# Swi4-dependent SWI4 transcription couples cell size to cell cycle commitment

**DOI:** 10.1101/2023.11.29.569332

**Authors:** Pooja Goswami, Abhishek Ghimire, Carleton Coffin, Jing Cheng, Jasmin Coulombe-Huntington, Ghada Ghazal, Yogitha Thattikota, Mike Tyers, Sylvain Tollis, Catherine Royer

## Abstract

Growth-dependent accumulation of the limiting SBF transcription factor, composed of Swi4 and Swi6, occurs in G1 phase in budding yeast and is limiting for commitment to division, termed Start. Here we measure size-dependence of Swi4 protein copy number under different genetic contexts using the scanning number and brightness technique. Mutation of SBF binding sites in the *SWI4* promoter or disruption of SBF activation resulted in ∼33-50% decrease in Swi4 accumulation rate and concordantly increased cell size at Start. Ectopic inducible expression of Swi4 in G1 phase cells increased production of Swi4 from the endogenous promoter, upregulated transcription of the G1/S regulon, and accelerated Start. Despite the potential for Swi4 positive feedback, G1 phase Swi4 accumulation was linear unless the Whi5 transcriptional repressor was inactivated. A threshold model in which Swi4 titrates SBF binding sites in G1/S promoters predicted the effects of nutrients, ploidy, and G1/S regulatory mutations on cell size. These results exemplify how transcription factor auto-production can contribute to a cell state transition.

## Introduction

The G1/S transition of the cell cycle is controlled at the transcriptional level in organisms from yeast to humans (*1*, *2*). In budding yeast, this transition (termed Start) is activated by the SBF and MBF transcription factor (TF) complexes (*3*), which govern expression of ∼200 genes in the G1/S regulon. These complexes comprise the same Swi6 activator subunit, and distinct DNA binding subunits, Swi4 and Mbp1, respectively. SBF and MBF bind to target sites, SCB and MCB, present in promoters of the G1/S regulon (*3–5*). The timing of the G1/S transition relative to cell growth determines cell size at Start. While deletion of *MBP1* in cells grown on rich nutrients does not significantly increase cell size, *swi4Δ* mutants are extremely large, demonstrating that under these conditions, SBF is the dominant transcriptional activator of Start (*6*). Transcriptional activation by SBF is repressed during G1 phase by Whi5 which binds to and inactivates SBF (*7*). The SBF-Whi5 complex is phosphorylated on multiple sites by the Cln-Cdc28 cyclin-dependent kinases in late G1, resulting in its dissociation and Whi5 export from the nucleus which is the earliest molecular indicator of Start (*8*, *9*). Deletion of the upstream G1 cyclin Cln3 strongly increases cell size, due to a Start delay with respect to growth (*10*). The triple *CLN1/2/3* deletion or inactivation of Cdc28 causes permanent arrest at Start (*11–13*). Importantly, *CLN1/2* genes are part of the G1/S regulon, and form a phosphorylation-driven positive feedback loop that activates SBF (*14*). Deletion of *WHI5* causes a small cell size phenotype that is epistatic to deletion of *SWI4* (*7*), underscoring the central role of SBF at Start.

We have shown previously by scanning Number and Brightness (sN&B) fluorescence microscopy that the absolute copy numbers of Swi4 and Mbp1 in early G1 phase are limiting with respect to their ∼200 target G1/S promoters (*6*). Copy numbers of all SBF/MBF subunits increase during G1 to levels that eventually saturate G1/S promoters. In particular, Swi4 copy number is much lower than that of Mbp1, Swi6 or Whi5 in small cells but increases ∼5-6-fold throughout G1 phase, leading to a doubling of its nuclear concentration. In contrast, absolute protein copy number accumulation of Swi6, Mbp1 and Whi5 scales with growth, yielding constant nuclear concentrations in G1 phase(*6*). The level of Whi5 concentration through G1 phase is controversial since relative measurements by epifluorescence microscopy suggest that Whi5 may be diluted as cells grow (*15*, *16*), although the apparent signal reduction may arise from photobleaching and other effects (*17*). The positive differential accumulation of Swi4 with respect to growth and to the other G1/S TFs raises the key question of how the expression of Swi4 during G1 phase is regulated. An upstream activating sequence (UAS) in the *SWI4* promoter contains several SCB/MCB sites, as well as an Early Cell Cycle Box (ECB) (*18*). The Mcm1 transcriptional activator, which binds the ECB (*19*), contributes to Swi4, Swi6 and Mbp1 production (*20*, *21*). Deletion of 140 base pairs corresponding to this UAS was shown to cause a 10-fold decrease in Swi4 levels (*18*). Deletion of *SWI6* also decreased Swi4 levels and abrogated its cyclic production (*18*), suggesting a contribution from the SCB/MCB sites to Swi4 expression. Furthermore, genome-wide CHIP-seq experiments demonstrated a physical interaction of Swi4 with its own promoter (*4*, *21*). Based on these observations, Swi4-dependent Swi4 expression has been postulated (*4*, *18*, *21*, *22*). However, this mechanism has never been quantitatively established, nor its potential role at Start evaluated. Moreover, it is unknown whether MBF contributes to Swi4 production.

In this study we quantitatively established the effects of Swi4, other G1/S regulatory proteins and transcriptional regulatory elements within the *SWI4* promoter on Swi4 production. Using the sN&B particle counting technique (6), we found that deletion or mutation of different combinations of the SCB/MCB sites in the *SWI4* promoter resulted in a ∼33-46% decrease in the rate of accumulation of Swi4 copy number with respect to growth. Disruption of phosphorylation of the SBF-Whi5 complex, but not *MBP1* deletion, resulted in a ∼50% decrease in Swi4 accumulation rate, further implicating SBF in its own production. Ectopic expression of Swi4 from a β-estradiol (BE2)-dependent promoter led to a -dependent increase of Swi4 expression from the endogenous promoter, which was accompanied by upregulation of genes in the G1/S regulon. A threshold of Swi4 copy number, in active SBF complexes, appeared to be required for Start and suggested that titration of G1/S promoters gates Start in various environmental and genetic contexts. Together, these results demonstrate that SBF-mediated *SWI4* expression contributes to the timing of the G1/S transition.

## Results

### Swi4 binding sites in the SWI4 promoter contribute to Swi4 protein expression

To test the role of Swi4 in its own production we used 2-photon (2p) sN&B particle counting to determine absolute values of intracellular concentrations of fluorescently tagged proteins (*6*). We measured the absolute nuclear concentration of Swi4 fused to fast folding-monomeric GFPmut3 (hereafter referred to as Swi4-GFP for brevity) expressed from its endogenous locus in single cells asynchronously growing in SC+2% glucose medium (*6*). Individual cell sizes were determined from the single cell images, and for simplicity the nuclear volume was assumed to be ∼1/7^th^ of the total cell volume (*23*), although we note that the scaling of nuclear volume with size in G1 is actually slightly sub-linear (*24*). Swi4 nuclear copy number was calculated as the product of the average nuclear concentration and the estimated nuclear volume. Since Swi4 exits the nucleus sometime after budding but before mitosis (*6*), the measured cells with nuclear Swi4 were either in G1 or S-phase.

We compared Swi4 protein copy numbers as a function of cell size in asynchronous populations of wild-type (WT) Swi4-GFP versus Swi4-GFP strains bearing various deletions or mutations in the *SWI4* promoter region (Fig. 1, S1, Table S1). In addition to three clustered central SCB/MCB promoter sites (termed site C in Fig. 1) situated ∼30 bp upstream of the ECB site documented previously (*18*), we identified two other putative SCB sites, one situated ∼1200 bp (termed site U) and the other ∼270 bp upstream of the *SWI4* ORF (termed site D) that could potentially contribute to *SWI4* expression.

**Figure 1.**
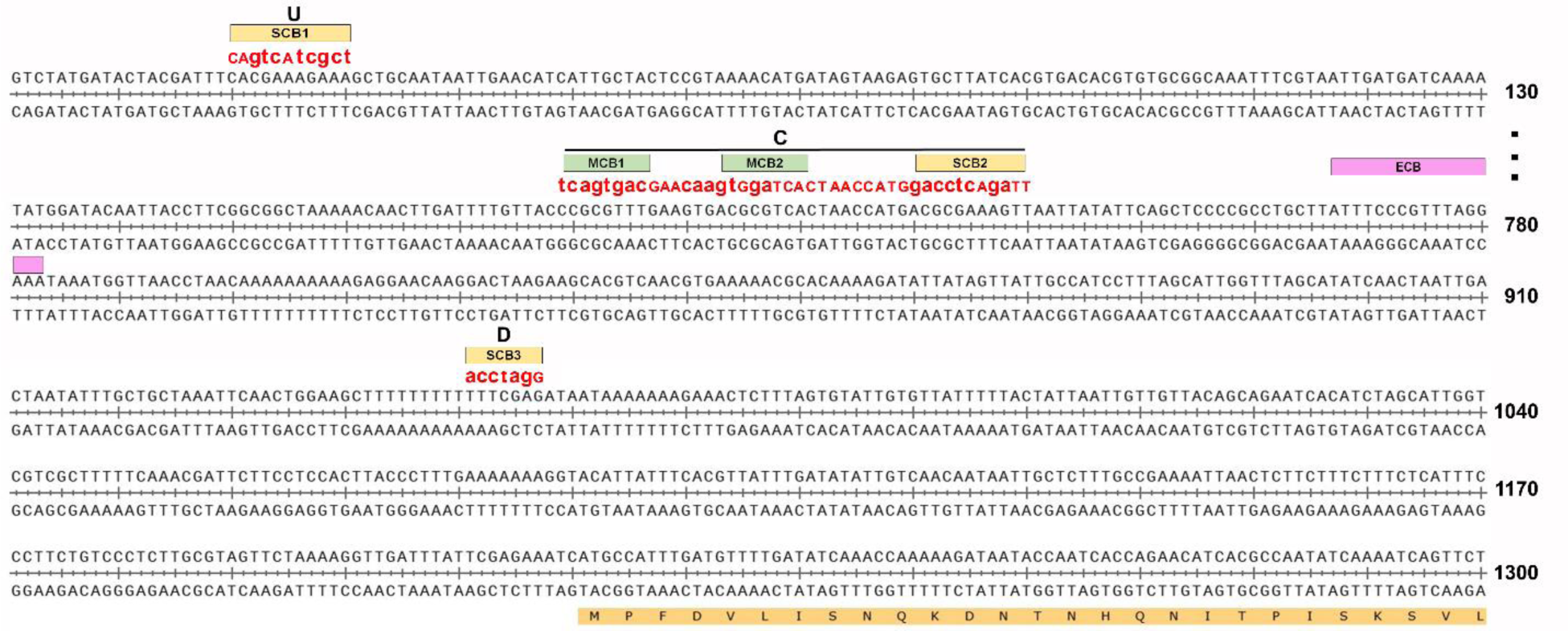
Mutations of the SCB/MCB sites in the *SWI4* promoter. Figure 1. Mutations of the SCB/MCB sites in the SWI4 promoter. Region of the *SWI4* promoter containing the SCB/MCB sites deleted or mutated in this study. Green rectangles designate MCB sites, orange rectangles designate SCB sites, pink rectangle designates the ECB site which was not deleted in our strains. U refers to the upstream SCB1 site, C to the central site containing MCB1, MCB2 and SCB2. D refers to the downstream SCB3 site. The beginning of the *SWI4* open reading frame (ORF) is also shown. WT and mutated sequences are listed in Table S1.

Deletion of the central region harboring two MCB sites and one SCB site (a total of 39 bp deleted, termed C delete, *P_SWI4Cdel_-SWI4-GFP*) decreased the rate of accumulation of Swi4-GFP protein copy number with respect to cell size (dSwi4/fL) by 33% compared to WT, while mutation of the same central sites to non-consensus SCB/MCB sequences (C mutations, *P_SWI4Cmut_-SWI4-GFP*) resulted in a slightly larger (37%) decrease (Fig. 2A, Table 1, Table S1). This same mutation of the central promoter region, together with mutation of the downstream SCB site (C+D mutations, *P_SWI4C+Dmut_-SWI4-GFP*) also decreased Swi4-GFP accumulation (∼38%), while additional mutation of the upstream SCB site (C+D+U mutations, *P_SWI4C+D+Umut_-SWI4-GFP*) decreased Swi4 accumulation even further (46%) (Fig. 2B, Table 1). These results establish that the MCB/SCB sites in the *SWI4* promoter are responsible collectively for about half of total Swi4 protein expression in G1-S phase haploid cells. Interestingly, in heterozygous *SWI4/SWI4-GFP* diploid cells (*i.e.*, WT promoter for both alleles but only one allele labeled with GFP), the rate of accumulation of labeled Swi4 nuclear copy number with respect to size was 58% lower than in haploids, while the deletion of the central promoter on the remaining allele expressing Swi4-GFP lowered the accumulation rate even further (67%, Table 1, Fig. S2). The total Swi4 accumulation rate with respect to size in heterozygous *SWI4/SWI4-GFP* diploids corresponded to only a 16% increase compared to haploids, even though there were two *SWI4* loci and twice as many G1/S TF target sites in diploids. We do not know the basis for this reduced Swi4 accumulation in diploids, but it may help explain why diploids are approximately twice as large as haploids.

**Figure 2.**
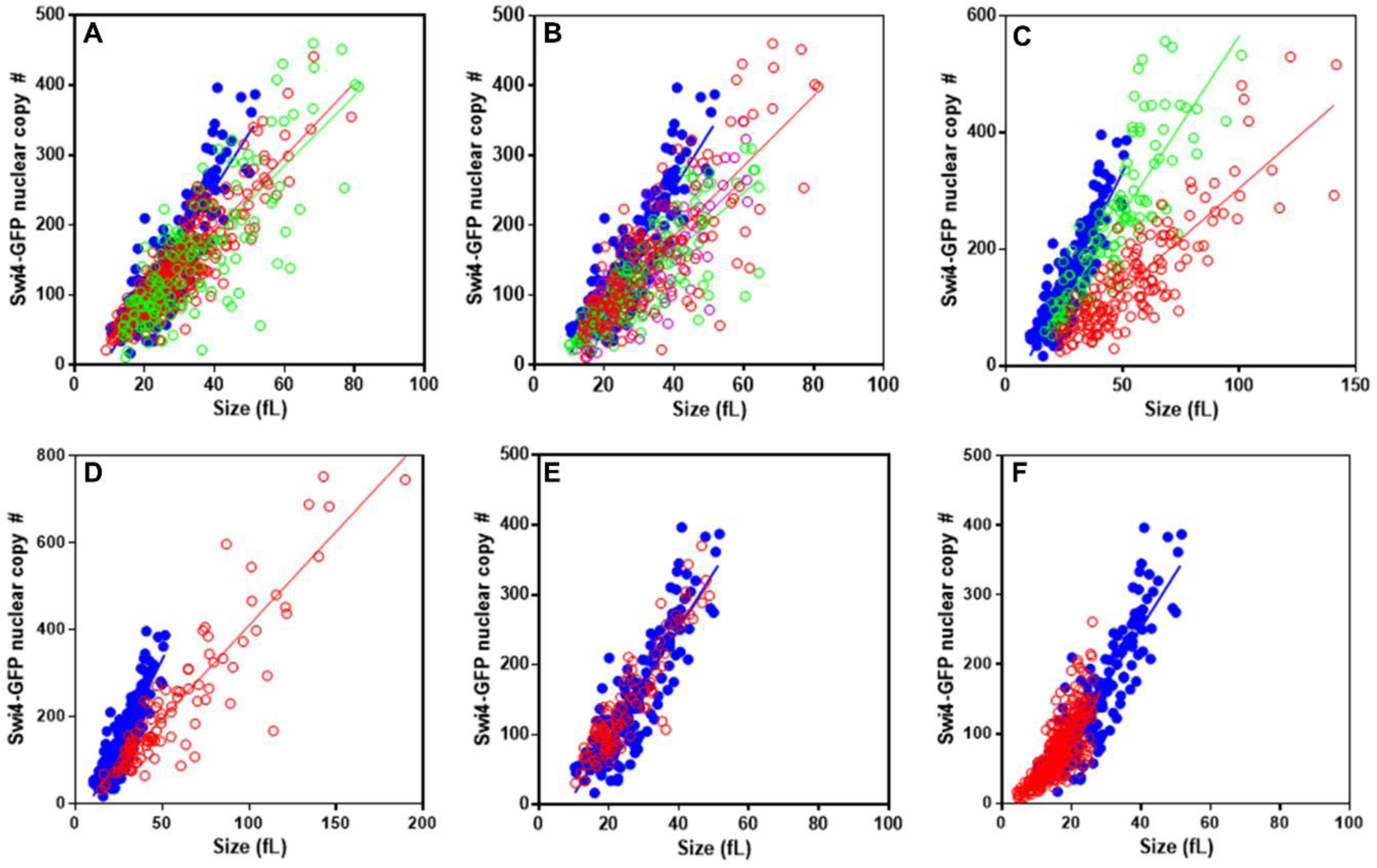
SCB/MCB elements in the Swi4 promoter regulate Swi4 protein production. A,B) SCB/MCB sites in the *SWI4* promoter contribute to the production of Swi4-GFP. Nuclear copy number of Swi4-GFP in (A) WT (blue), C deletion, *P_Cdel_-SWI4-GFP* (red) and C mutation, *P_Cmut_-SWI4-GFP* (green) strains; (B) WT (blue), C mutation, *P_Cmut_-SWI4-GFP* (red), C+D mutations, *P_C+Dmut_-SWI4-GFP* (pink) and C+D+U mutations, *P_C+D+Umut_-SWI4-GFP* (green) strains; (C) WT (blue), *cdc28as1* in absence of 1-NM-PP-1 (green) and *cdc28as1* in presence of 10 μM 1-NM-PP-1 (red); D) WT (blue) and *cln3Δ SWI4-GFP* (red) and E) WT (blue) and *mbp1Δ SWI4-GFP* (red); F) WT (blue) and *whi5Δ SWI4-GFP* (red). Number of cells were WT Swi4-GFP (MTy5270) 115, *P_Cdel_-SWI4-GFP* (MTy5320) 387, *P_Cmut_-SWI4-GFP* (MTy5314) 142, *P_C+Dmut_-SWI4-GFP* (MTy5298) (130), *P_C+D+Umut_-SWI4-GFP* (MTy5302) (138), *cdc28as1 SWI4*-*GFP* (MTy5317) in absence of 1-NM-PP-1 116, and in presence of 1-NM-PP-1 160, *cln3Δ* Swi4-GFP (MTy5318) 113, *mbp1Δ* Swi4-GFP (MTy5271) 101, and *whi5Δ* Swi4-GFP (MTy5180) 297.

**Table 1.**
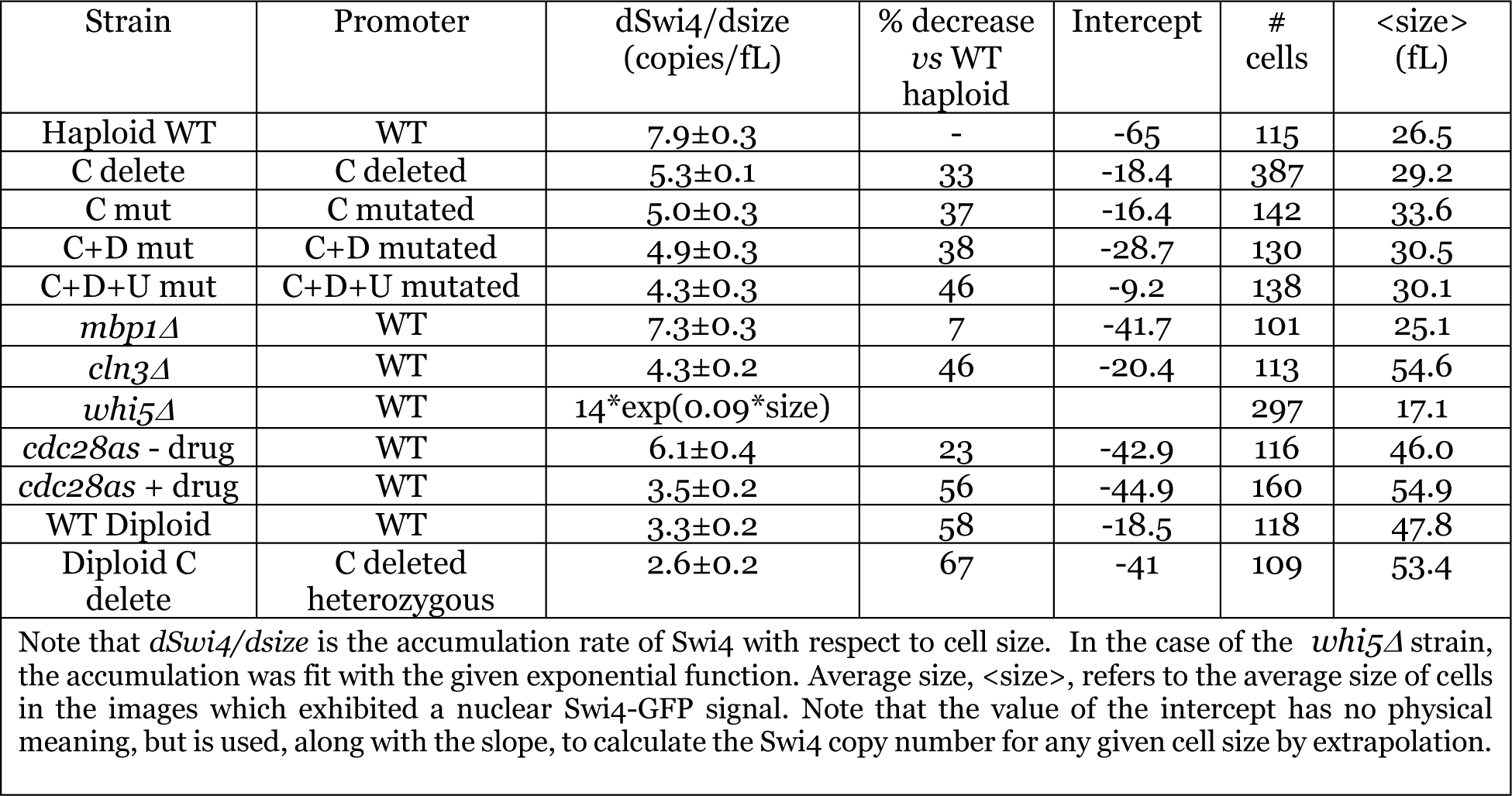
Effect of mutations and deletions on Swi4-GFP accumulation vs cell size.

In agreement with our previously published model, in which Swi4 dosage is a limiting factor for cell cycle commitment (*6*), deletions or mutations of the various MCB/SCB sites in the *SWI4* promoter in haploid cells led to a modest large size phenotype, as assessed by Coulter counter measurements (Fig. S1, S3A-C, S4, Table S2). The average size, as derived from the images of single cells exhibiting nuclear Swi4-GFP, hence in G1-S phase, was also larger for the promoter mutants compared to WT cells (Fig. S1, Table 2). This phenotype was reduced in *SWI4-GFP* heterozygous diploids, where the mean size for the heterozygous diploid in which one allele carried the *P_SWI4Cdel_*-*SWI4-GFP* was ∼2fL larger than WT diploid (Fig. S3, see also Table S2, below). These results showed that mutations that diminish the rate of Swi4 production led to a delay in Start, and a larger cell size.

**Table 2.**
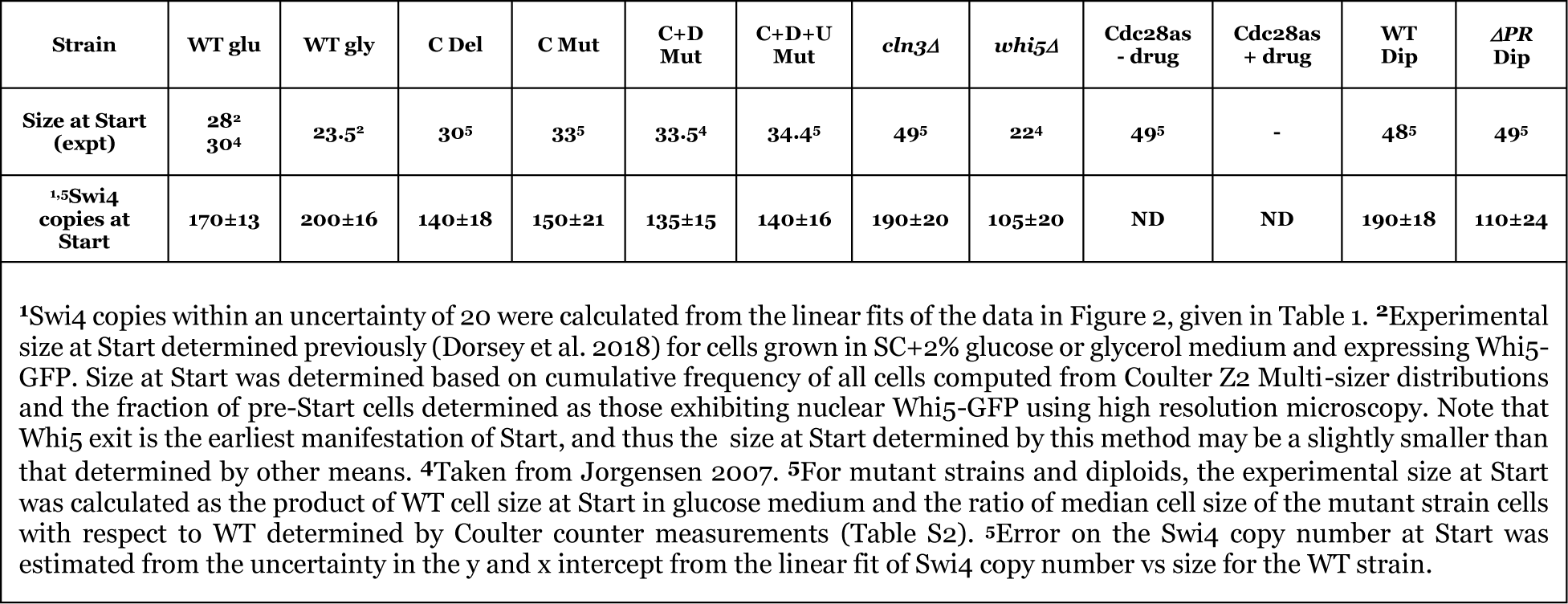
Predicted Swi4 copy number at Start.

### Cln3-Cdc28 mediated phosphorylation is required for proper Swi4 accumulation

The dependence of Swi4 expression on SCB/MCB sites in the *SWI4* promoter indicated that SBF and/or MBF are involved in Swi4 production. Since SBF activation requires Cdc28 activity, we evaluated the Cdc28-dependence of Swi4 accumulation. Chemical inhibition of kinase activity in a strain expressing an analog sensitive Cdc28, *cdc28-as1* (*25*), with 10 μM of the inhibitor, 1-(tert-butyl)-3-(naphthalene-1-methyl)-1H-pyrazolo [3,4-d] pyrimidine-4-amine (1-NM-PP-1) resulted in a large decrease in Swi4-GFP copy number accumulation (56%) with respect to cell size (Fig. 2C, Table 1). Hence, Cdc28 kinase activity drives approximately half of Swi4 accumulation in G1 phase. The rate of Swi4-GFP accumulation in absence of the inhibitor was slightly lower (23%) than for WT cells, perhaps due to impaired kinase activity of the Cdc28-as1 protein. We verified the requirement for G1 phase Cdc28 activity by removing Cln3, which activates Cdc28 in early G1 phase. *CLN3* deletion decreased the rate of accumulation of Swi4-GFP as a function of cell size by ∼46%, similar to the decrease observed upon mutating all MCB and SCB sites (Fig. 2D, Table 1). The large size phenotype of the *CDC28-as1* and *cln3Δ* strains was apparent in the large average cell sizes observed for cells exhibiting nuclear Swi4-GFP in our images (Fig S1, Table 1) and in Coulter counter measurements (Fig. S3F, Tables 1, S3). Taken together, these results demonstrate that Cln3-Cdc28 kinase activity contributes to half of Swi4 accumulation in G1, suggesting that SBF activation is critical for Swi4 accumulation.

Due to the functional overlap between SBF and MBF, and the shared ability of Swi4 and Mbp1 to bind the same SCB/MCB sites (*4*, *5*, *22*, *26*), it is in principle possible that MBF could contribute to Swi4 expression. However, the lack of any effect of *MBP1* deletion on the rate of Swi4 copy number accumulation versus cell size (Table 1, Fig. 2E), suggested Mbp1 does not mediate Swi4 production. Swi4 accumulation in a strain deleted for the repressor of SBF, Whi5, led to non-linear accumulation with respect to size (Fig. 2F). In a *whi5Δ* strain, the size dependence of Swi4 copy number was best described with an exponential function (Table 1). Newborn daughter *whi5Δ* cells are very small, 5 fL, and had only ∼14 copies of Swi4 (∼7 dimers), which is extremely sub-stoichiometric with respect to the ∼200 G1/S promoters. We note that the cell-to-cell variation in Swi4 concentration for all strains examined was much larger than the variation in Swi4 protein copy number (Fig. S5). Since copy number is computed as the nuclear Swi4 concentration multiplied by the nuclear volume, this means that the stochasticity is mostly associated with the growth rates of individual cells, and not the Swi4 accumulation rate, which is more robust.

### Ectopic Swi4 protein expression increases endogenous Swi4 protein levels

A prediction of Swi4-dependent *SWI4* expression is that an acute ectopic increase of Swi4 in G1 cells should stimulate endogenous *SWI4* transcription and an increase in endogenous Swi4 protein. We therefore examined the effect of ectopic expression of unlabeled Swi4 on the production of Swi4-GFP from the endogenous promoter. We constructed a strain that constitutively expresses a chimeric Z3EV transcription factor from the *ACT1* promoter with an additional copy of the *SWI4* gene under the control of a *Z3EV*-responsive promoter at the *BUD9* locus (*27–29*). The Z3EV transcription factor is composed of the human Znf268 zinc-finger DNA binding domain, which recognizes a 9 base pair sequence, fused to the ligand binding domain of the estrogen receptor, which is responsive to β-estradiol (BE2). Hence, the activity of this chimeric TF, and therefore the transcription of the exogenous *SWI4* gene cassette, is BE2-dependent.

In a control strain to assess BE2-dependent expression, Swi4-GFP was expressed from the *Z3EV* promoter in the cassette, while the endogenous copy of *SWI4* remained unlabeled (*P_Z3EV_-SWI4-GFP SWI4*). We used this strain to measure how much Swi4 protein was produced by the *Z3EV* promoter as a function of BE2 concentration. BE2-dependent expression and nuclear localization of Swi4-GFP was observed as expected, with intensities of Swi4-GFP at a background level in absence of (Fig. 3A, Fig. S6) and reaching saturation in presence of 5 nM BE2. We note that prolonged overexpression of Swi4-GFP in this strain upon addition of 5 nM BE2 (Fig. 3A) led to cytokinesis defects (Fig. S6D).

**Figure 3.**
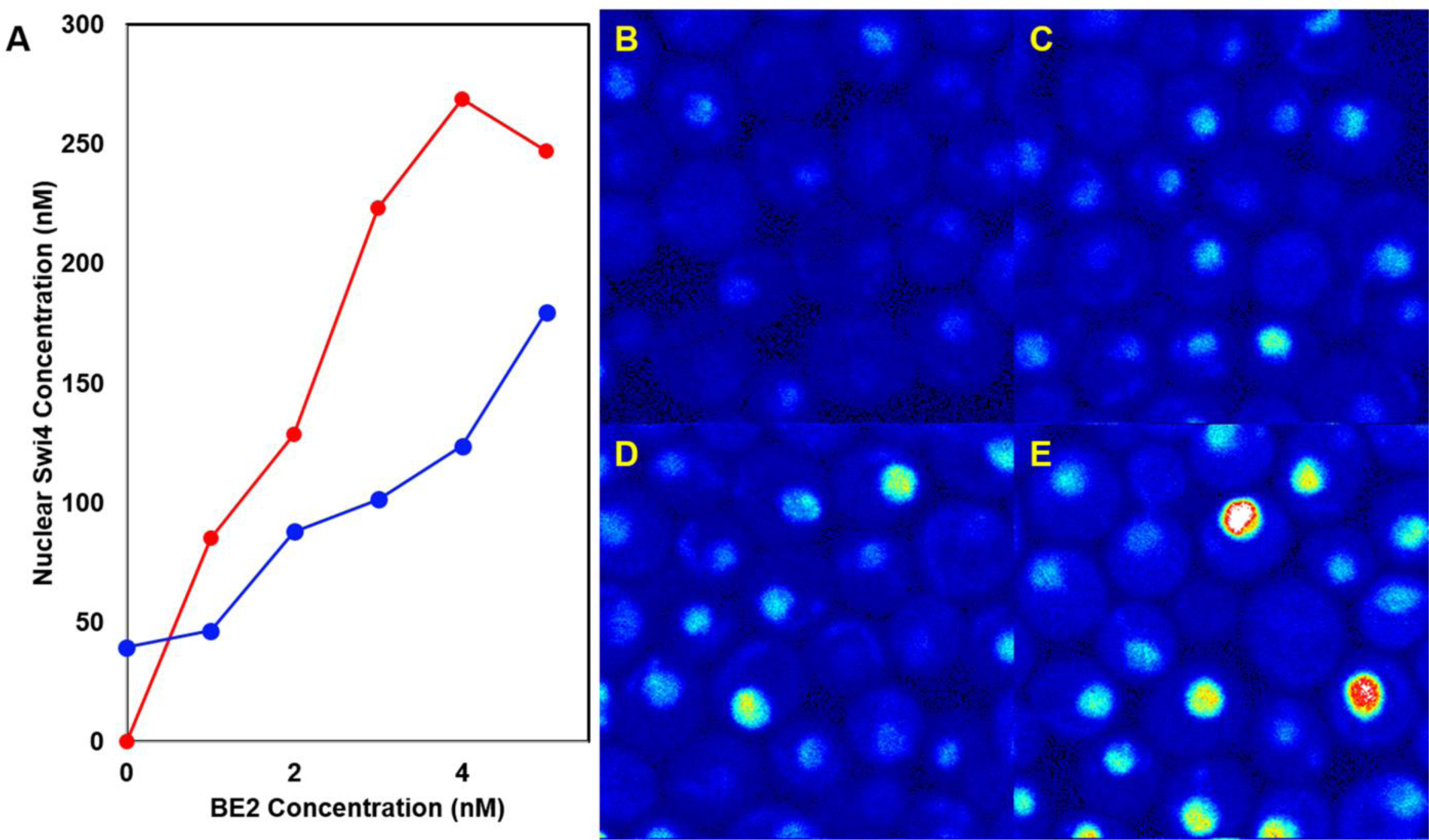
BE2-dependent production of Swi4-GFP. A) Swi4-GFP nuclear concentration as a function of BE2 concentration for the *P_Z3EV_-SWI4 SWI4-GFP* strain (MTy5189) (blue) and the control *P_Z3EV_-SWI4-GFP SWI4* strain (MTy5190) (red). B-E) Images of FOV of the *P_Z3EV_-SWI4 SWI4-GFP* strain (MTy5189) in presence of 0, 2, 3, and 5 nM BE2, respectively. Image full scale is 0-8 counts. Note that in absence of BE2, the average nuclear intensity was higher in the *P_Z3EV_-SWI4 SWI4-GFP* strain than for the control strain due to the presence of endogenous Swi4-GFP (*6*). FOV are 20×20 µm.

In an experimental strain, the exogenous Swi4 expressed from the *Z3EV* promoter was unlabeled, while Swi4-GFP was expressed from the endogenous *SWI4* promoter *(P_Z3EV_-SWI4 SWI4-GFP*). We used this strain to measure how much Swi4 protein was produced by the endogenous promoter in response to ectopic Swi4 induction as a function of BE2 concentration (Fig. 3A-blue, B-D). We observed a substantial increase in the endogenous Swi4-GFP expression with increasing concentrations of BE2. This result confirmed a key prediction, *i.e*., that additional ectopic Swi4 should increase endogenous Swi4-GFP expression.

As over-expression of Swi4-GFP led to a cytokinesis defect in asynchronous populations, homeostatic cell size at the population level could not be used as a readout for acceleration of Start in response to BE2 in the Z3EV strains. To directly examine the effect of Swi4 over-expression on the timing of Start we collected small G1 daughter cells from the *P_Z3EV_-SWI4 SWI4-GFP* strain synchronized by centrifugal elutriation. We then measured the time-dependence of Swi4-GFP production in absence of BE2 and in presence of 5 nM BE2 using imaging conditions that minimized photo-bleaching (*17*). In presence of 5 nM BE2, the endogenous Swi4-GFP intensity increased markedly as a function of time, whereas in the absence of the signal was comparable to that observed in the WT strain expressing Swi4-GFP (Fig. 4A, B).

**Figure 4.**
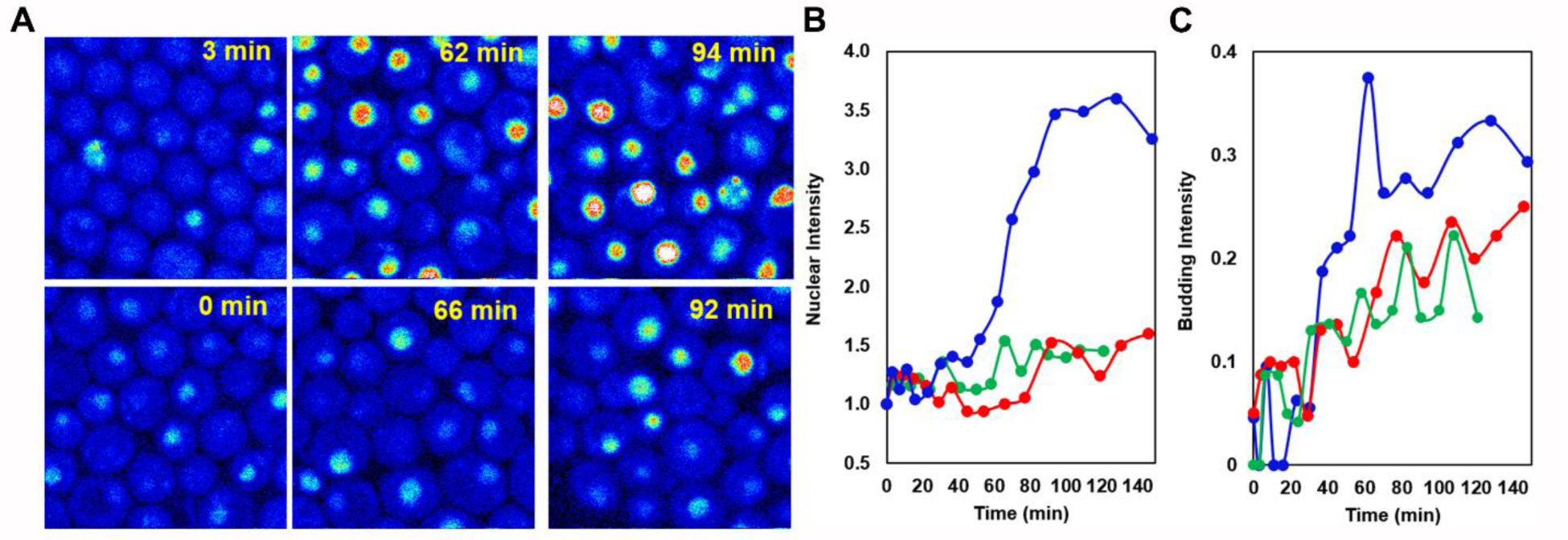
Effect of BE2-dependent exogenous Swi4 production on *SWI4* transcription. A) Images of fields of view (FOV) of elutriated G1 daughter cells from the *P_Z3EV_-SWI4, SWI4-GFP* strain (MTy5189) in presence of 5 nM BE2 (top row) and in absence of BE2 (bottom row), as a function of time after addition of BE2 to the cells in the top row. Time points are indicated in the images. Intensity scale is 0 (black) to 2 (white) photon counts per 40 μs pixel dwell-time. FOV are 20×20 µm. B) Normalized intensity of nuclear Swi4-GFP as a function of time for the *P_Z3EV_-SWI4 SWI4-GFP* strain (MTy5189) in presence of 5 nM BE2 (blue), in absence of BE2 (red) and the WT Swi4-GFP strain (green). C) Budding index as a function of time for the *P_Z3EV_-SWI4 SWI4-GFP* strain (MTy5189) strain in presence of 5 nM BE2 (blue), in absence of BE2 (red) and the WT Swi4-GFP strain (green). Cells were grown in SC+2% glucose medium. Average number of cells quantified per time point was 22 ±2 for the WT Swi4-GFP time-course, 19 ±2 for the *P_Z3EV_-SWI4 SWI4-GFP* strain (MTy5189) in presence of 5 nM BE2 and 20 ±2 in absence of BE2.

### Increasing Swi4 levels accelerates the timing of Start

We have previously shown that Swi6 copy number in G1 phase is in excess compared to Swi4 and, moreover, that the Swi4/Whi5 ratio in WT cells grown in glucose medium depends on cell size, increasing from ∼0.42 in small cells to ∼0.84 in large G1 cells (*6*). We thus expected that increased Swi4 production in response to would increase the number of active SBF complexes to levels sufficient to overcome Whi5 inhibition, leading to an acceleration of Start. In time course imaging experiments, we found that the budding index, i.e. the fraction of cells in the fields of view (FOV) exhibiting buds, increased more rapidly in presence of 5 nM BE2 than in its absence for the *P_Z3EV_-SWI4 SWI4-GFP* strain (Fig. 4C).

Premature budding should also correlate with an increased expression of the G1/S regulon as ectopic Swi4 accumulates. To test this hypothesis, we performed a whole-genome RNA-sequencing time-course following treatment of a *P_Z3EV_*-*SWI4 SWI4* strain in absence and presence of 5 nM BE2. BE2-dependent expression of *SWI4* led to an increase in expression of several key G1/S regulon genes, including *CLN1* and *PCL1* (Fig. 5A,B, Table S3A). A number of SBF and MBF target genes were also down-regulated in a time dependent manner after addition, including a known MBF target, *CDC21* (Fig. 5C,D, Table S3B). These results demonstrate that ectopic production of Swi4 is sufficient to prematurely activate G1/S transcription and Start.

**Figure 5.**
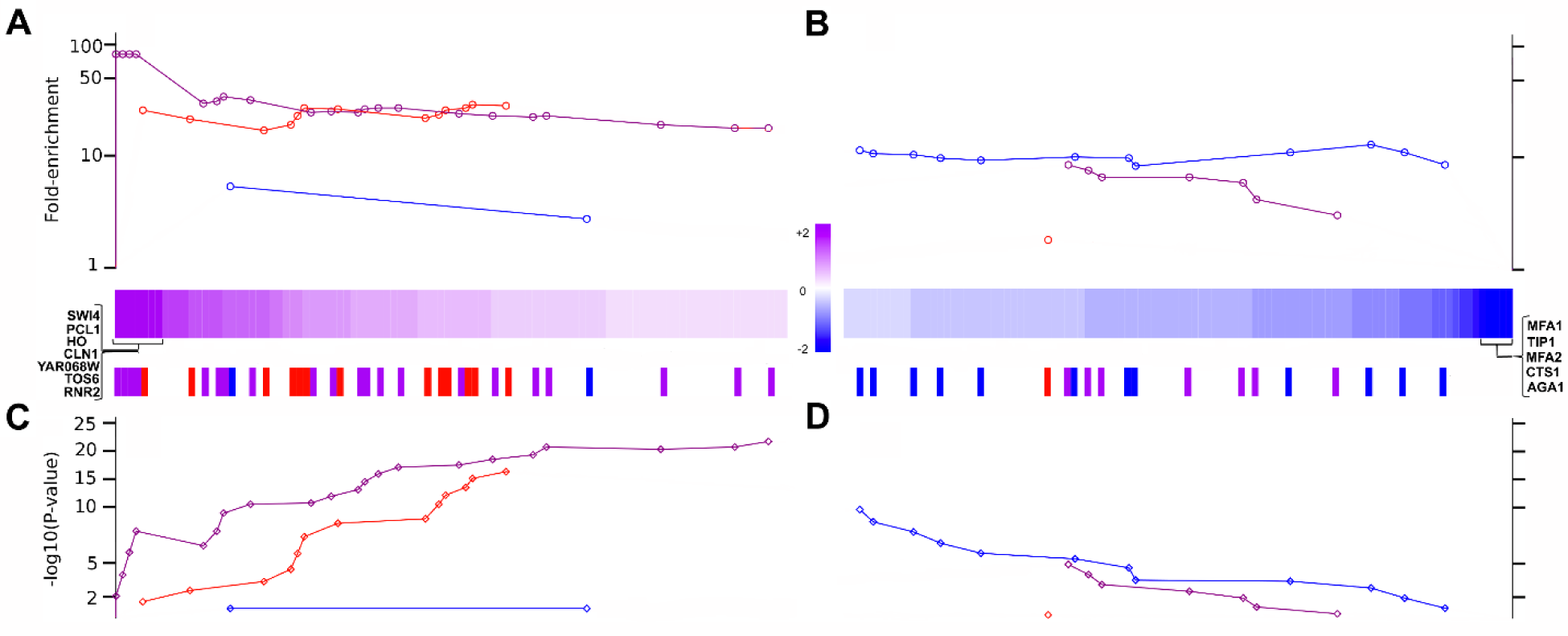
Effect of BE2-dependent exogenous Swi4 production on transcription of the G1/S regulon. A and B) Fold enrichment (up-regulation) and fold repression (down-regulation) in presence of 5 nM BE2. Colored strips represent genes that were up-regulated (pink) and downregulated (blue). The seven most strongly up-regulated genes are shown on the left, while the five most strongly down-regulated genes are shown on the right. Red bars below the fold enrichment strips corresponds to SBF target genes, blue bars to MBF target genes and purple bars to both (41). C and D) –logP value for up-regulation (C) and down-regulation (D) compared to untreated and ethanol treated cells. Only genes with an absolute log2-fold change are shown. In panel B lack of up-regulation was normalized to a fold-enrichment of 1, while the expression of the most down-regulated genes in panel D was normalized to 1. A complete list of the detected up- and down-regulated genes and quantitative values is provided in Table S2A,B. Strain used was *P_Z3EV_-SWI4 SWI4* (MTy5188).

### A threshold of Swi4 protein copy number gates Start

The above results revealed that genetic contexts that affect the critical cell size at Start also modulate the rate of Swi4 accumulation with respect to growth rate, suggesting the existence of a potential threshold in Swi4 copy number required for Start. To test this idea we computed the number of Swi4 molecules present in WT cells grown in glucose medium when they pass Start, and compared this number to the Swi4 copy number that corresponds to the estimated critical cell size at Start in other genetic/environmental conditions, given their distinct Swi4 accumulation rates with respect to growth. We previously measured that WT cells are 28 fL at Start based on the cumulative fraction of Whi5-GFP-containing WT cells grown in the same conditions (*6*), in reasonable agreement with the size observed for onset of budding, 30 fL (*23*). From the Swi4-GFP copy number vs cell size plot (Figure 2A), we estimated that WT cells grown in glucose have accumulated about 170 (+/-13) Swi4 molecules when they reach 28 fL and Start (Table 2).

Using the Swi4 accumulation rate versus cell size across genetic backgrounds (Table 1), we computed the number of Swi4 copies at Start for other strains and conditions. The size at Start was estimated from the ratio of the median daughter cell size obtained by Coulter counter measurements of the mutant strains to that of the WT strain (Table S2), multiplied by the size at Start for the WT strain (28 fL). The promoter mutant strains were slightly larger than WT, with a correspondingly larger estimated size at Start (Table 2). The corresponding number of Swi4 copies present in these cells at Start was slightly lower than for WT cells, but was within the uncertainty as deduced from the linear fit of the plot of WT Swi4 copy number vs size (Figure 2). For diploid cells, based on the measured Swi4 accumulation rate (Table 1), 190 (+/-18) Swi4 molecules per genome were predicted at the estimated size at Start of 48 fL. This Swi4 copy number at Start value was also fairly close to that found for haploid cells (170). Thus, the lower Swi4 accumulation rate per haploid genome in diploid cells (Table 1), whatever its origin, accounts reasonably well for the relationship between size and ploidy. Likewise, using our prior results for Swi4 copy number accumulation vs size for WT haploid cells grown in glycerol (*6*), a Swi4 copy number of 200±16 predicted a size at Start of 23±2 fL. While the threshold is slightly higher than in glycerol, the predicted size is quite close to the previously measured value of 23.5 fL (*6*). From this simple analysis we conclude that a threshold of 170-200 Swi4 molecules per haploid genome appears to gate Start across those strains and conditions.

This threshold of ∼170-200 Swi4 copies was determined above for strains in which the phosphorylation module of the G1/S regulatory network is intact. We next calculated the Swi4 copy number present at Start for the *cln3Δ* strain in which SBF-Whi5 phosphorylation is compromised. The experimental size at Start for the *cln3Δ* strain, estimated as above (Table 2, (*30*)), is 49 fL. Based on the linear fit for Swi4 accumulation in this strain (Table 1), the Swi4 copy number at Start was 190 (±20), similar to the estimated threshold value for WT. Note that the proportion of cells in G1 phase is ∼7% larger in the *cln3Δ* strain relative to WT (*31*), such that the size at Start, and thus, the corresponding Swi4 copy number, could be 7% larger than 190 (∼200), still in reasonable agreement with the WT threshold. In the *whi5Δ* strain, cells passed Start at a smaller size, estimated at 22 fL (Table S2, (*7*)). This size corresponded to a substantially lower Swi4 copy number, 105 (±20), based on the experimentally observed supra-linear relationship between Swi4 copy number and cell size (Figure 2, Table 1). Hence, *whi5Δ* cells, in which all SBF complexes are de-repressed, pass Start with significantly less Swi4 than WT cells, suggesting that fewer, but more active, SBF complexes may reduce the threshold requirement for Swi4 copy number at Start.

## Discussion

The role of Swi4-dependent *SWI4* transcription in the Start transition is supported by four main lines of evidence: (*i*) quantitative sN&B measurements show that the rate of accumulation of Swi4 versus size depends on the presence of Swi4 target sites in the *SWI4* promoter; (*ii*) dependence of Swi4 accumulation rate on Cln3-Cdc28 activity; (*iii*) ectopic Swi4-mediated induction of endogenous Swi4 expression and acceleration of G1/S transcription; (*iv*) decreased rate of Swi4 accumulation in diploids with respect to growth and concomitantly larger cell size. Swi4-dependent *SWI4* transcription is responsible for approximately half of Swi4 accumulation in G1 phase in WT haploid cells grown in glucose medium. Interestingly, the contribution of Swi4 autoregulation to Swi4 accumulation corresponds to the difference in accumulation rate of Swi4 with respect to that of the other G1/S TFs (Swi6, Whi5, and Mbp1), all of which scale approximately linearly with growth and exhibit size-independent concentrations (*6*). The interconnected feedback loops of Swi4 production, Cln1/2 production (*14*) and SBF/Whi5 phosphorylation (*8*, *9*, *32*) all likely contribute to a size-resolved, sharp and effectively irreversible Start transition.

Our measurements suggest that in wild type cells, a threshold of ∼170 Swi4 monomers (∼85 SBF complexes, see (*6*)) is required to activate Start. 85 SBF complexes would titrate on average slightly less than half of the ∼200 promoters in the G1/S regulon. This Swi4 threshold accounts reasonably well for the difference in cell size for diploids compared to haploids and the effects of mutations in the *SWI4* promoter. This threshold also rationalizes the small size at Start of WT cells grown in poor nutrients, due to a positive differential scaling of Swi4 accumulation with respect to growth. From these observations we suggest that titration of a threshold fraction of G1/S promoters appears to be required for Start. This promoter titration by Swi4 hypothesis is reminiscent of a previous titration model in which G1/S promoters are titrated by Cln3 (*33*).

Titration of G1/S promoters, while necessary, is not sufficient to trigger Start, as evidenced by the behavior of the analog sensitive *cdc28as1* strain in presence of 1NM-PP-1 inhibitor (*25*), which accumulates Swi4 to WT levels, yet never passes Start. Thus, the Start transition requires not only that a given fraction of the G1/S promoters be bound by SBF, but also that some fraction of the bound SBF complexes are *active*. Hence, promoter titration by SBF and SBF/Whi5 phosphorylation combine to achieve a threshold number of *active* SBF complexes required to pass Start. A decrease in phosphorylation activity by mutation (e.g., in a *cln3Δ* strain) results in a decreased rate of Swi4 accumulation with respect to size, and therefore, an increase in cell size at the required threshold of Swi4 molecules. Apparently, in this case, sufficient phosphorylation activity by Cln1/2-Cdc28 complexes is achieved when *cln3Δ* cells attain the WT Swi4 copy number threshold (170) to activate the fraction of SBF required for Start.

In contrast, in absence of Whi5 repression, cells pass Start at a smaller size (*7*) and at a lower Swi4 threshold than WT cells because SBF is not inhibited by Whi5. The number of Swi4 molecules at Start in *whi5Δ* cells (80–100) may correspond to the threshold of active SBF complexes in WT cells. We note, however, that Swi6 copy number is slightly limiting in cells of all sizes with respect to the sum of Swi4 and Mbp1 copies (*6*). Moreover, there may be a residual requirement for Swi6 phosphorylation, even in absence of Whi5, to achieve complete activation of SBF. Indeed, cells that over-express Cln3 are smaller than *whi5Δ* cells (*34*, *35*). Interestingly, the inherent positive feedback in Swi4 production appears to be unmasked in the *whi5Δ* strain. While WT Whi5 levels repress most SBF copies and therefore make the amount of active SBF only weakly dependent on the amount of available Swi4, in absence of Whi5 more Swi4 molecules immediately lead to more active SBF copies and therefore to an *increase in the rate* of Swi4 accumulation, yielding a supra-linear (possibly exponential) increase of Swi4 copy number with cell size. This observation suggests that in WT cells, the phosphorylation requirement hinders Swi4 positive feedback.

While we have clearly observed that differential scaling of Swi4 accumulation with respect to growth modulates cell size at Start, the molecular mechanisms responsible for this phenomenon remain to be determined. For example, why does the Swi4 accumulation rate slow less than bulk growth rate in poor nutrients? And similarly, while the observed lower Swi4 accumulation rate in diploids may account for the size relationships of ploidy, the molecular mechanism for this reduced accumulation rate per haploid genome in diploids remains to be discovered. Overall, the linked Swi4 accumulation/phosphorylation feedback loops provide distinct molecular inputs for modulation of cell size in response to mutations or growth conditions, and combined, lead to a sharp G1/S transition. Irreversible switch-like transitions are implicated in many important cellular processes such as development, differentiation and damage responses. Transcription factor autoregulation and titration of target promoters in complex regulons, such as that reported here, may be a recurrent regulatory motif in many of these transitions.

## Star Methods

### Experimental Design

The objective of the study was to quantify the effects of key G1/S factors on the accumulation of the Swi4 transcription factor during cell growth in G1. To this purpose, we fused GFPmut3 to the Swi4 C-terminus at the endogenous *SWI4* locus, and quantified the Swi4-GFP protein levels using the scanning Number and Brightness particle counting approach as implemented in (*6*). Those measurements were performed in several genetic backgrounds bearing deletions/mutations of key genes or their promoters known to control the G1/S transition. Budding yeast strains used in this work are summarized in Table 3. None of the GFP-tagged experimental strains exhibited noticeable cell size and growth phenotype difference from their untagged counterparts.

**Table 3.**
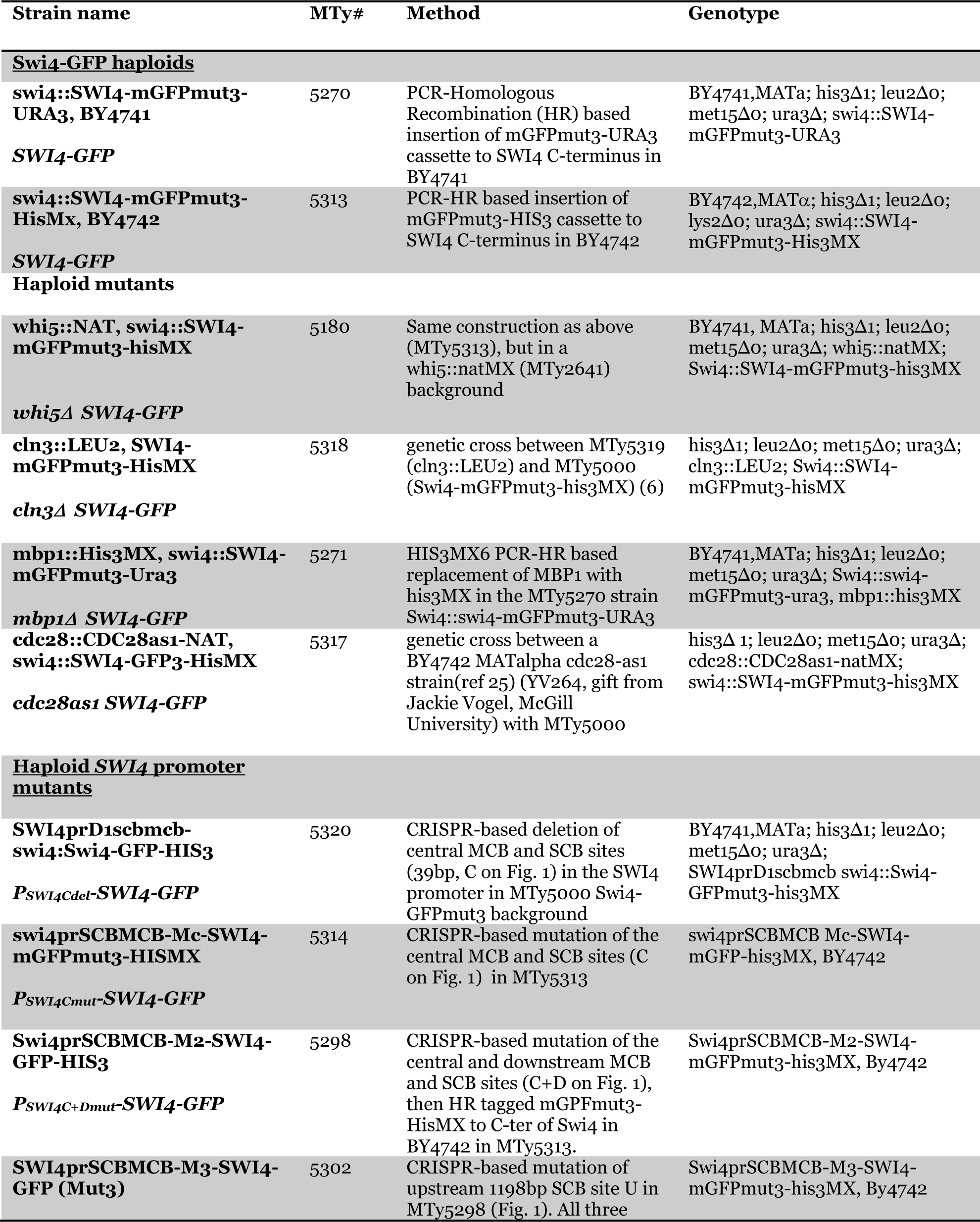
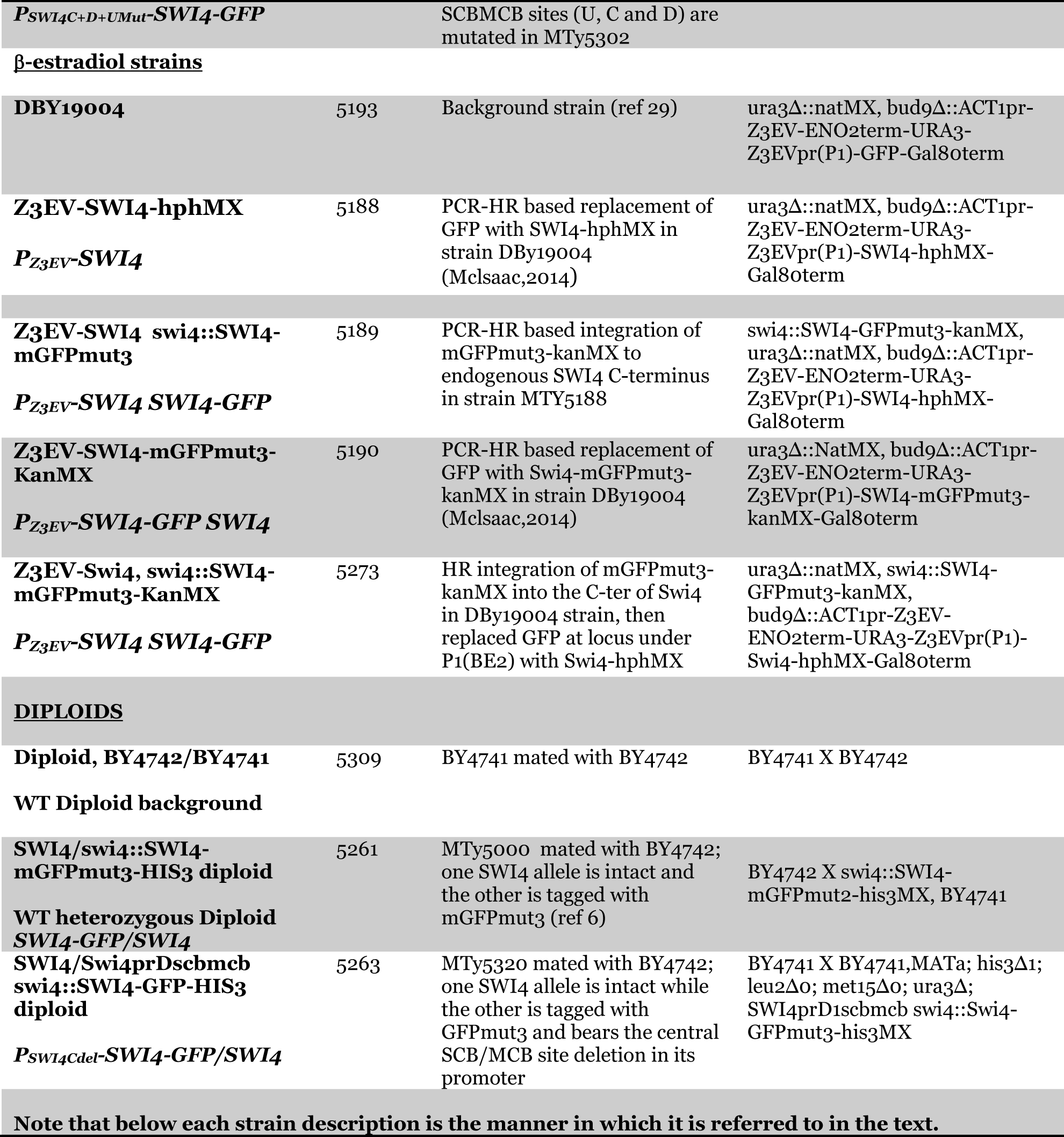
Strains Used.

### Yeast Strain Construction

All haploid experimental strains were generated in the BY4741 or BY4742 derivatives of the S288C background as indicated, with PCR-based homologous recombination integration of a *mGFPmut3-his3MX* cassette (or corresponding *URA3* or *kanMX* versions) at the C terminus of the Swi4 natural locus, which were crossed into other BY4741 mutant strains as indicated. A strain producing monomeric GFP from an inducible promoter, used in molecular brightness calculations, was obtained by transforming plasmid into BY4741 strain. This strain is referred to as “free GFP strain” in the manuscript.

The β-Estradiol (BE2) inducible strains were constructed by using Z3EV expression strain DBy19004 (MTy5193) (*36*) together with integration of the mGFPmut3-HIS3MX cassette at the endogenous *SWI4* locus, using ura3Δ::natMX bud9Δ::ACT1pr-Z3EV-ENO2term-URA3-Z3EVpr(P1)-GFP-Gal80term as the background strain. In this strain, the Z3EV artificial transcription factor controlled by the ACT1 promoter and ENO2 terminator was integrated to replace the non-essential *BUD9* locus to express Z3EV and any gene of interest under the Z3EV-responsive promoter Z3EVpr(P1) (*P_Z3EV_)*, which contains binding sites for Z3EV and allows conditional expression upon β-estradiol addition to the culture. The strain referred to as *P_Z3EV_-SWI4 SWI4-GFP* with unlabeled Swi4 expressed from the Z3EV promoter and *swi4::SWI4-GFPmut3* (MTy5189) was used to assess Swi4-dependent *SWI4-GFP* induction. Note GFPmut3 is referred to in the following and elsewhere in the text as GFP. PCR fragment based homologous recombination was used to integrate *SWI4-hphMX* under the Z3EV promoter (P1) to replace GFP in strain DBy19004 and yield the *P_Z3EV_-SWI4* strain MTy5188. The mGFPmut3-KanMX cassette was then integrated at endogenous *SWI4* locus to obtain the strain *P_Z3EV_-SWI4 SWI4::SWI4-GFP* (MTy5189). A control strain, *P_Z3EV-_SWI4-GFP SWI4* (MTy5190), was constructed in the same manner to demonstrate Swi4-GFP production from the *Z3EV* promoter and measure induction at different BE2 concentrations.

*SWI4* promoter deletions and mutations were introduced by CRISPR/Cas9 mediated replaced with mutated repair sequences. Briefly, guide RNAs (gRNAs) were cloned into plasmid pGZ110 (kindly provided by Bruce Futcher, SUNY Stonybrook) and co-transformed with a SWI4 promoter repair cassette (either cloned into pGZ110 or as a separate linear fragment) to introduce the mutations of interest. gRNA and repair cassette sequences are shown in Table 4. Target strains contained either wild type *SWI4* allele or a *SWI4-mGFPmut3* cassette integrated at the endogenous locus. All mutations were confirmed by sequencing of genomic DNA. As indicated in Table 3, promoter mutations were combined with other strain genotypes by genetic crosses.

**Table 4.**
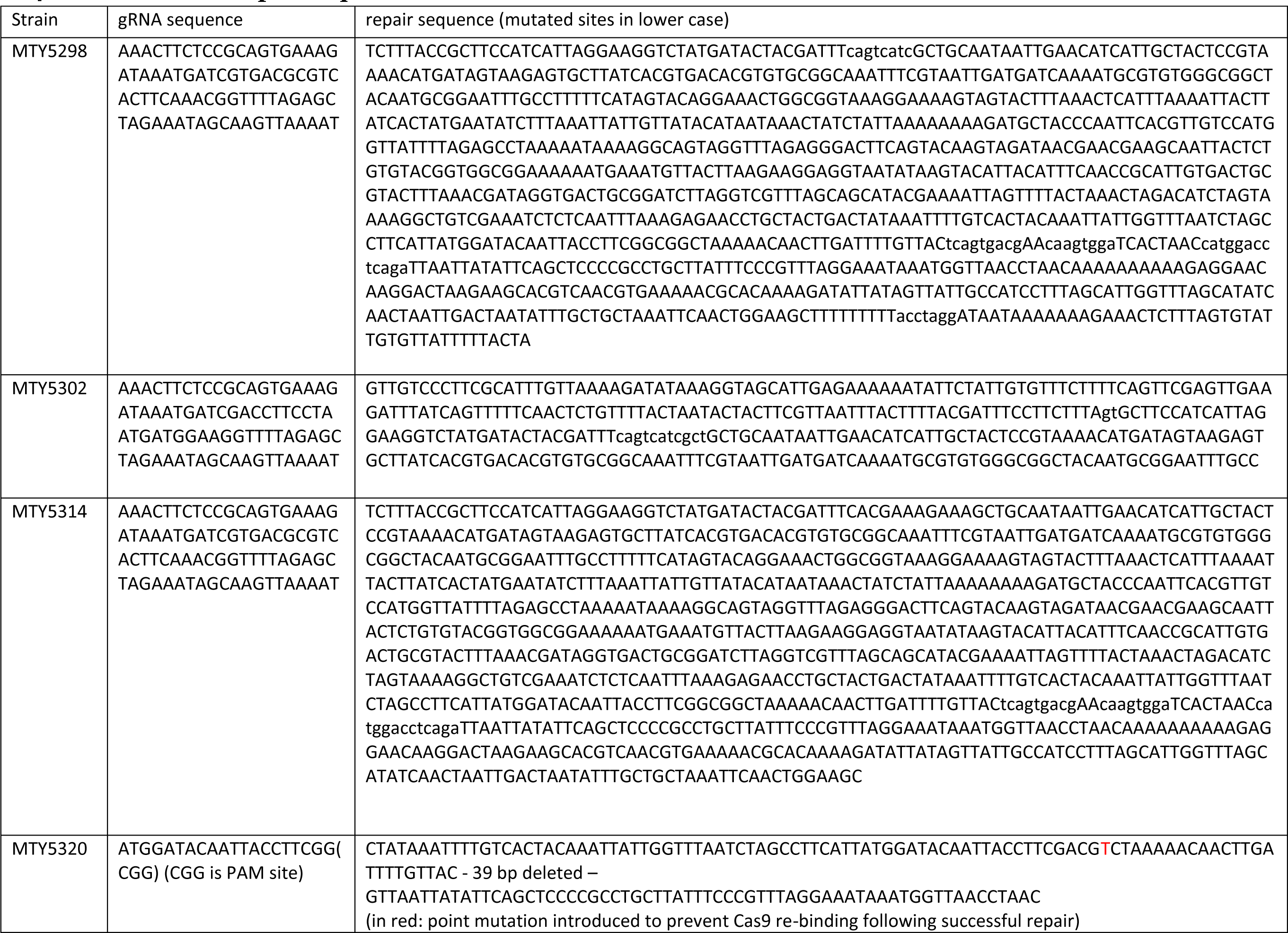
Guide RNA and Repair sequences used in strain construction.

### Cell Culture

All strains were grown in Synthetic Complete (SC) medium with 2.0 g/L amino acid mix (Sunrise Science) supplemented with 6.7 g/L yeast nitrogen base (Sigma-Aldrich) and 2% w/v glucose. For each sN&B experiment, a single colony of appropriate (untagged) background strain, free GFP strain and the GFP-tagged strains of interest were grown in 1 mL SC 2% glucose medium to saturation overnight. The saturated overnight cultures were diluted 1/30 in 1 mL SC 2% glucose medium and grown to early exponential phase (4-5 h) before imaging. For synchronous time course experiments in presence of BE2, early G1 cells obtained by centrifugal elutriation were induced with 5 nM or 0 nM BE2 in SC 2% glucose media for 15 min at 30^0^C, and added to SC 2% glucose agarose pads containing 5 nM or 0 nM BE2 prior to imaging. The strain expressing the analog sensitive allele of the Cdc28 kinase was induced with 10 μM ATP analog drug; 1-(tert-butyl)-3-(naphthalen-1-ylmethyl)-1H-pyrazolo [3,4-d] pyrimidin-4-amine, referred to as 1-NM-PP-1, directly on agarose pads. Cells were grown on these pads for 20 minutes prior to imaging. For GFP induction of the GAL1-GFPmut3 strain used for GFP brightness calibration (free GFP strain), after overnight growth to saturation in SC glucose medium, the culture was washed, diluted 1/30 and grown in SC 2% raffinose medium for 3-4 h, followed by the addition of 0.1% galactose to the culture for 1 h. The induced culture was then washed twice with SC 2% glucose medium and grown in SC 2% glucose medium for 3-4 h before imaging under similar growth and culture conditions as the control and protein fusion strains.

### Preparation of agarose pads

Before each sN&B experiment, 200 µL from a 1 mL culture grown to early exponential phase (OD600 of 0.16-0.18, cell density 1-2 × 10^6^ cells/mL) was pelleted in a microfuge at 1300 rpm for 30 seconds and 195 µL of the supernatant was discarded. Cells were immediately re-suspended in the remaining 5 µL medium, and 3.5 µL of the suspension was mounted onto a 2% glucose agarose pad (62µL) as previously described (Dorsey et al, 2018). The agarose pad containing mounted cells was sealed with a ConA (Sigma, 2 mg/mL)-coated coverslip for immobilization of cells during imaging. The sealed pads were clamped in an Attofluor chamber (Molecular Probes) and clusters of monolayer cells were imaged for no longer than 2 h.

### Elutriation of BE2 dependent strains

Centrifugal elutriation was used to isolate synchronous G1 cells for the time-course imaging of WT Swi4 (MTy 5270) and BE2 strain (MTy 5189). Cells were cultured in 1.2 L of SC + 2% glucose overnight to early exponential growth phase (0D_600_=0.14 measured using a Tecan Infinite M1000 Pro Plate Reader), and centrifuged in a Sorvall SLC-6000 rotor at 1100 rpm for 10 min at 4°C. Cells were resuspended in 50mL of the supernatant, lightly sonicated (30 s, 1 s intervals at power level 1), loaded in a Beckman JE-5.0 rotor at 8 mL/min at 1500 rpm and then eluted at 12 mL/min in a 50 mL fraction. 200 µL of the elutriated G1 fraction was pelleted in a microfuge at 1300 RPM for 30 seconds and 195 µL of the supernatant was discarded. Cells were immediately re-suspended in the remaining 5 µL medium. Finally, 3.5 µL of the suspension was mounted onto a 2% glucose agarose pad (62µL) as previously described (*6*).

### Scanning Number and Brightness (sN&B) imaging

For accurate quantitative measurement of absolute concentrations of Swi4, we used an image based particle-counting fluctuation microscopy technique called sN&B (*37*). The sN&B acquisitions were performed in an ISS Alba fast scanning mirror fluctuation microscope (ISS, Champaign, IL) equipped with 2-photon laser excitation (Mai Tai Ti: Sapphire, Newport-SpectraPhysics, Mountain View, CA) at an excitation wavelength of 1000 nm. Emission from GFP was obtained using a 530 +/- 50 nm band-pass filter. Acquisition for each strain was done in multiple Fields of View (FOVs) with the FOV size of 256×256 pixels covering 20×20 μm. Each FOV was scanned with 50 raster scans at a 40 μs pixel dwell-time. Further, each FOV was imaged in 3 z-positions separated by 500 nm.

### Steady-state and time course imaging of BE2-dependent strains

In steady-state BE2 experiments, the WT Swi4-GFP strain (MTy 5270) and the experimental and control *Z3EV* strains, MTy5189 and MTy5190, respectively, were imaged as for all other strains except that in addition cells were incubated for 30 minutes in SC medium containing 0-5 nM of BE2 after reaching the early exponential phase of growth. The BE2-induced cultures were centrifuged, the supernatant removed, and induced cells were grown in 1mL SC 2% glucose media for 1 h before mounting on agarose pads for imaging.

In the BE2 time courses, early G1 cells of WT Swi4-GFP (MTy 5270) and the BE2 strains (MTy 5189 and MTy5190) obtained through centrifugal elutriation were treated with 5 nM BE2 for 15 minutes and mounted onto a 2% glucose agarose pad as described above. The cells were imaged as above at 31% power. To avoid photobleaching, a new FOV was imaged at every time point for 10 scans at 40 µs pixel dwell time at 3 z-positions separated by 500 nm.

### Scanning Number and Brightness Image Analysis

Methodology for sN&B analysis has been described previously (Dorsey et al., 2018) but is briefly summarized here. For absolute quantification of fluorescent proteins using sN&B, we used probabilistic auto-fluorescent background subtraction, based on comparing the distribution of background intensities from hundreds of millions of pixels of images from cells that do not express GFP fusions in order to generate a probabilistic calculation of the auto-fluorescent background intensity contribution at every pixel in the FOV from cells expressing specific GFP fusions. Two calibration parameters were performed for obtaining the absolute concentrations of GFP-tagged proteins. First, the effective volume of the microscope, *V_PSF_*, defined by laser alignment, was obtained for each experiment by sN&B using a solution of 46 or 28 nM fluorescein in Tris buffer (pH 8.0) and 40% glycerol. Second, an *in cellulo* sN&B calibration of the molecular brightness of free monomeric GFP, *e_GFP_*, was performed using cells expressing monomeric GFP from a plasmid (*6*) to compute the concentrations of the GFP fusion proteins. For the Swi4-GFP strains examined, the absolute concentration of nuclear Swi4-GFP in each cell was calculated from the average intensity of all nuclear pixels after background subtraction, *<F>_nuc_*, as:

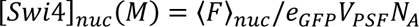

where *N*_A_ is the Avogadro number. Nuclear volume for individual cells was calculated as 1/7 of the total cell size (*23*) derived from whole cell masking and assuming a sphere.

All sN&B acquisitions were analyzed using custom 2015 MATLAB scripts (Sylvain Tollis, University of Montreal with edits from Steven Notley, RPI), which has previously been used to determine absolute concentration of fluorescent proteins in live cells (*6*, *38*). This semi-automated analysis allowed the computation of absolute protein concentrations in the best Z-position for selected regions of interest i.e., the nucleus of pre-Start cells.

### Intensity Analysis

All analysis of fluorescence intensity of Z3EV strains were performed in the analysis software SimFCS (Enrico Gratton, Laboratory of Fluorescence Dynamics, UC Irvine). Fluorescence intensity from 5 FOVs at each concentration were averaged and plotted in Figure 4. All fluorescence intensity images used SimFCS at 0-14 intensity units as the scale.

### RNA-seq

The *P_Z3EV_-SWI4 SWI4 (MTy5188)* cells were grown as described above for sN&B experiments and induced for 1 h with either 5 nM of BE2 (in ethanol solvent) or with ethanol as a negative control. Total RNA was prepared as described in (*39*) both before (pre-induction sample) and after induction. Briefly, cells were disrupted using glass beads, and their aqueous content was extracted using phenol/chloroform. Nucleic acids were then precipitated using ethanol and genomic DNA was removed using DNase treatment (Qiagen). RNA was quantified using Qubit (Thermo Scientific) and assessed for quality with a 2100 Bioanalyzer (Agilent Technologies). Poly-A selected transcriptome libraries were generated using the Kapa RNA HyperPrep (Roche). Sequencing was performed on an Illumina NextSeq 500 system at the IRIC genomic platform. Experiments were performed in duplicate.

RNA-sequencing data analysis was performed as detailed in (*38*). Briefly, raw reads were aligned to all Ensembl yeast transcripts using Bowtie 2.2.5 (*40*) with default parameters and counts defined as alignments with an edit distance smaller than 5. Dubious ORFs as annotated in the Saccharomyces Genome Database (yeastgenome.org) were excluded. Only genes with at least 500 reads in one sample were further analyzed. Gene expression levels were expressed as log2 read counts, normalized to the upper-quartile gene expression level for each sample separately (*41*). For each of the two replicates, the expression after 1 h induction was normalized separately to the expression in the pre-induction sample and in the control ethanol-treated sample of the same replicate, yielding four fold-change values for each gene. Then, the smallest of those log2 fold-change (in absolute value) was used to generate the final differential expression profile represented on Figure 5. Hence, a gene had to be upregulated (or downregulated) with respect to both the pre-induction *and* the ethanol-treated control sample across both replicates to yield a positive (or negative) score. The former control reduced sample-to-sample variation while the second eliminated BE2-independent but time-dependent gene expression variation in the cell population. Genes for which any of the four normalizations yielded log2 fold changes of different signs were assigned a score of 0. Among the most up- and down-regulated genes, direct SBF and/or MBF targets were identified based on the G1/S regulon gene list defined by Ferrezuelo et al. (*42*). At each ranking position n, the enrichment of SBF targets (for upregulated genes) or MBF targets (for downregulated genes) within the n^th^ top differentially regulated genes as compared to random expectations was computed, and the significance of the enrichment level was calculated using Fisher’s exact test.

### Coulter Counter Cell Sizing and Growth Curve

The population size distributions of all strains were determined using a Beckman Coulter Counter Z2 particle sizer calibrated to a default aperture size of 50 µm and 31.02 diameter calibration constant, with the upper and lower size threshold for particles set to 7.256 and 2.255 µm, respectively. Overnight cultures from a single colony were diluted 1:30 in 1 mL SC 2% glucose medium and grown to early exponential phase (4-5 hours). 200 µL of culture was added to 10 mL of Beckman Isoton II diluent solution and sonicated lightly for 30 seconds before size acquisition. The number of cells per binned cell size window were divided by the total particle count to yield the fraction of cells in each size bin, referred to as “cell size distribution” in the manuscript. For assessing the growth rate of different strains, cells were grown and diluted as above in SC 2% glucose medium and measured every hour in a Grenier 96-well flat well transparent plate at 600 nm (OD600). The doubling time was assessed from the slope of the linear part of the growth curve, that corresponds to log phase.

## Acknowledgments

This work was supported by grants Canadian Institutes of Health Research FDN-167277 to MT, the Academy of Finland (grant #350887) and the Sigrid Jusélius Foundation (grant #220196) to ST and NSF PHY 1806638 to CAR. We thank Bruce Futcher for kindly providing plasmid pGZ110, and Brenda Andrews, Jackie Vogel and Scott McIsaac for generously providing yeast strains.

## Author contributions

Investigation: PG, AG, CC, JC, GG, YT, ST Formal analysis: PG, AG, CC, JCH Resources. JC, GG, ST Data curation: JCH, CAR, AG Methodology: CAR, ST, MT Validation GG, JC, ST Conceptualization: ST, MT, CAR Software: ST, CAR, JCH Writing – original draft: ST, MT, CAR, AG, JC, JCH Writing – review & editing: ST, MT, CAR Project administration: ST, MT, CAR Funding acquisition: ST, MT, CAR

## Data access

All data needed to evaluate the conclusions in the paper are present in the paper and/or the Supplementary Materials. Imaging data will be made available upon request.

All strains will be provided by MT and ST pending scientific review and a completed material transfer agreement. Requests for the strains should be submitted to.

