## Supplementary material for "Swi4-dependent SWI4 transcription couples cell size to cell cycle commitment": Supplemary material

| Target | | | | | | | | | | | | | |
|---|---|---|---|---|---|---|---|---|---|---|---|---|---|
| SCB1 | | | C | A | C | G | A | A | A | G | A | A | A |
| SCB1mut | | | C | A | g | t | c | A | t | c | g | c | t |
| Consensus | | | C | R | C | G | A | A | | | | | |
| SCB2 | | A | C | G | C | G | A | A | A | G | T | T | |
| SCB2mut | | g | a | c | C | t | c | A | g | a | T | T | |
| Consensus | | | C | R | C | G | A | A | | | | | |
| SCB3 | T | T | T | T | C | G | A | G | A | | | | |
| SCB3mut | T | a | c | c | t | a | g | g | A | | | | |
| Consensus | | | C | R | C | G | A | A | | | | | |
| MCB1 | | C | C | C | G | C | G | T | T | T | | | |
| MCB1mut | | C | t | C | a | g | t | g | a | c | | | |
| Consensus | | | A | C | G | C | G | T | N | A | | | |
| MCB2 | T | G | A | C | G | C | G | T | C | A | | | |
| MCB2mut | a | a | g | t | G | g | a | T | C | A | | | |
| Consensus | | | A | C | G | C | G | T | N | A | | | |

**Table S1. SCB/MCB target sites in the *SWI4* promoter region**. Also given are their consensus sequences and the mutations used in the present study. In yellow highlight are bases that were mutated in the mutant strains while non-mutated bases are highlighted in peach.

**Table S2. Cell size mean and median from Coulter counter measurements and doubling time from growth curves**

| Strain | Median Cell Size 1 (fL) | Mean Cell Size 1 (fL) | Median Cell Size 2 (fL) | Mean Cell Size 2 (fL) | $\tau_D$ (min) |
|---|---|---|---|---|---|
| *BY4741 BG* | 53.02 | 61.21 | 51.65 | 59.51 | 108 |
| *WT SWI4GFP* | 55.20 | 63.89 | 56.40 | 65.19 | 105 |
| *$P_{Cmut}$-SWI4GFP* | 60.48 | 72.19 | 60.67 | 72.06 | 105 |
| *$P_{C+Dmut}$-SWI4GFP* | 60.05 | 71.79 | 59.21 | 71.38 | 97.6 |
| *$P_{C+D+Umut}$-SWI4GFP* | 61.55 | 72.88 | 69.49 | 69.49 | 105 |
| *$P_{Cdel}$-SWI4GFP* | 56.99 | 66.17 | 57.48 | 67.37 | 105 |
| *cln3Δ SWI4GFP* | 97.00 | 107.2 | 100.8 | 109.9 | 108 |
| *whi5Δ SWI4GFP* | 44.05 | 51.56 | 43.69 | 50.60 | 104 |
| *CDC28as1 SWI4GFP* | 89.14 | 99.47 | 90.34 | 102.0 | 102 |
| *Diploid BG* | 90.75 | 101.0 | 88.80 | 98.05 | 102 |
| *Diploid WT SWI4GFP* | 88.02 | 98.61 | 88.43 | 98.22 | 107 |
| *Diploid WTSwi4, $P_{Cdel}$-SWI4GFP* | 89.64 | 99.42 | 90.01 | 99.80 | 102 |
| *$P_{Z3EV}$_Swi4 SWI4GFP* | 66.50 | 76.73 | 68.27 | 77.35 | |
| *$P_{Z3EV}$_SWI4GFP SWI4* | 65.69 | 76.02 | 68.58 | 78.93 | |
| Average doubling time, $\tau_D$ | | | | | 105 ± 4 |

**Table S3A. RNA-sequencing results for genes that were up-regulated by .**

| Gene Name | Target of SBF (S), MBF (M), both (B), neither (N)* | Fold up-regulation |
|---|---|---|
| SWI4 | B | 5.67 |
| PCL1 | B | 2.97 |
| HO | B | 2.66 |
| TIR1 | N | 2.33 |
| CLN1 | B | 2.23 |
| TOS6 | S | 1.95 |
| YAR068W | N | 1.95 |
| RNR2 | N | 1.94 |
| RTC3 | N | 1.53 |
| URA3 | N | 1.49 |
| NFG1 | N | 1.48 |
| ICY1 | N | 1.45 |
| CSI2 | S | 1.44 |
| CWP1 | N | 1.4 |
| CLB1 | B | 1.37 |
| YHR214W | N | 1.36 |
| YHP1 | B | 1.32 |
| MNN1 | B | 1.29 |
| CLB5 | M | 1.27 |
| PUT4 | N | 1.26 |
| NRT1 | N | 1.24 |
| NRM1 | B | 1.23 |
| SWH1 | N | 1.18 |
| SUR1 | S | 1.18 |
| YAR066W | N | 1.15 |
| PBI2 | N | 1.11 |
| TGL5 | N | 1 |
| SVS1 | S | 0.98 |
| TOS7 | S | 0.96 |
| CLN2 | S | 0.87 |
| EXG1 | B | 0.86 |
| SRL1 | B | 0.85 |
| TMC1 | N | 0.85 |
| GAT2 | N | 0.85 |
| PRY2 | S | 0.78 |
| YPL088W | N | 0.77 |
| ATP20 | N | 0.75 |

| | | |
|---|---|---|
| GIC2 | B | 0.74 |
| RGL1 | N | 0.71 |
| YBR071W | B | 0.71 |
| YOX1 | B | 0.68 |
| AIM20 | N | 0.66 |
| IZH4 | N | 0.66 |
| CWP2 | B | 0.66 |
| PHD1 | N | 0.65 |
| YNL058C | N | 0.64 |
| PCL2 | S | 0.61 |
| PIG2 | N | 0.61 |
| FKS3 | N | 0.6 |
| KCH1 | S | 0.59 |
| SKM1 | S | 0.57 |
| CRH1 | B | 0.55 |
| YBR085C-A | N | 0.55 |
| CLB2 | S | 0.54 |
| MSB2 | S | 0.53 |
| PYC1 | N | 0.52 |
| DAP1 | N | 0.5 |
| YPL162C | N | 0.49 |
| HCM1 | B | 0.48 |
| PRB1 | N | 0.47 |
| RAX2 | S | 0.45 |
| MSF1 | N | 0.45 |
| HOR7 | N | 0.44 |
| NCW1 | N | 0.44 |
| WSC4 | N | 0.43 |
| YOR342C | B | 0.43 |
| CIK1 | N | 0.43 |
| XKS1 | N | 0.4 |
| COX9 | N | 0.39 |
| OCH1 | B | 0.39 |
| YDR061W | N | 0.39 |
| YLR345W | N | 0.38 |
| GNA1 | N | 0.38 |
| VVS1 | N | 0.37 |
| AFR1 | N | 0.37 |
| RSM18 | N | 0.36 |
| AXL2 | M | 0.36 |
| MXR1 | N | 0.36 |
| MRPL11 | N | 0.35 |
| BBP1 | N | 0.34 |

| | | |
|---|---|---|
| BIO2 | N | 0.34 |
| COX13 | N | 0.34 |
| TOS3 | N | 0.34 |
| SRL3 | N | 0.33 |
| SDC1 | N | 0.33 |
| SKS1 | N | 0.33 |
| YUH1 | N | 0.32 |
| MET14 | N | 0.32 |
| RNR1 | B | 0.32 |
| RTS3 | N | 0.32 |
| SNC2 | N | 0.31 |
| GPI11 | N | 0.31 |
| ARF2 | N | 0.3 |
| IZH3 | N | 0.29 |
| SPC42 | N | 0.29 |
| MSB3 | N | 0.29 |
| YET1 | N | 0.29 |
| HHT2 | B | 0.28 |
| SUT2 | N | 0.28 |
| RSM10 | N | 0.28 |

**Table S3B. RNA-sequencing results for genes that were down-regulated by .**

| Gene Name | Target of SBF (S), MBF (M), both (B), neither (N)* | Fold down-regulation |
|---|---|---|
| MFA1 | N | -4.45 |
| TIP1 | N | -2.19 |
| MFA2 | N | -1.95 |
| CTS1 | N | -1.9 |
| AGA1 | N | -1.75 |
| FRE4 | N | -1.48 |
| YLL053C | N | -1.41 |
| PHO3 | N | -1.41 |
| AI1 | N | -1.37 |
| DSE2 | N | -1.35 |
| SIT1 | N | -1.19 |
| CDC21 | M | -1.16 |
| MCM7 | N | -1.16 |
| PHO89 | N | -1.08 |
| DSE1 | N | -1.03 |
| AGA2 | N | -0.95 |
| CDC9 | N | -0.94 |
| YRO2 | N | -0.94 |
| MSA1 | M | -0.91 |
| MSA2 | M | -0.88 |
| STE2 | N | -0.82 |
| YGP1 | N | -0.82 |
| YBL113C | N | -0.81 |
| MCM3 | N | -0.81 |
| GAS3 | M | -0.81 |
| FIT2 | N | -0.8 |
| ARN1 | N | -0.77 |
| SPI1 | N | -0.76 |
| YLL067C | N | -0.75 |
| YHR219W | N | -0.73 |
| YML133C | B | -0.72 |
| SCW11 | N | -0.72 |
| FRE7 | N | -0.71 |
| MCM5 | N | -0.71 |
| WHI5 | N | -0.7 |
| YJL225C | N | -0.7 |
| ISR1 | N | -0.69 |

| | | |
|---|---|---|
| RBH2 | M | -0.66 |
| UTH1 | N | -0.66 |
| ATO3 | N | -0.66 |
| YEL077C | N | -0.65 |
| HSP30 | N | -0.65 |
| YIL177C | N | -0.64 |
| IRC7 | N | -0.64 |
| AI2 | N | -0.63 |
| YRF1-3 | B | -0.63 |
| ENB1 | N | -0.62 |
| YRF1-7 | B | -0.61 |
| FET3 | N | -0.61 |
| YRF1-2 | N | -0.6 |
| LYS20 | N | -0.59 |
| YLL066C | N | -0.59 |
| YRF1-4 | N | -0.59 |
| MCM4 | N | -0.58 |
| YRF1-5 | B | -0.56 |
| OPT1 | N | -0.56 |
| FCY21 | N | -0.56 |
| ARG3 | N | -0.55 |
| YRF1-8 | N | -0.54 |
| ARG1 | N | -0.54 |
| CIN2 | N | -0.53 |
| LYS2 | N | -0.53 |
| RFA1 | M | -0.53 |
| POL30 | N | -0.53 |
| KCC4 | N | -0.52 |
| YPR204W | N | -0.52 |
| RAD27 | M | -0.52 |
| PHO5 | N | -0.51 |
| YRF1-6 | B | -0.51 |
| YHL050C | N | -0.49 |
| ALG14 | B | -0.49 |
| ASF1 | M | -0.48 |
| RNH201 | N | -0.48 |
| YRF1-1 | B | -0.48 |
| ZRT1 | N | -0.47 |
| PRY1 | S | -0.47 |
| HMS2 | N | -0.47 |
| LYS9 | N | -0.45 |
| MPS3 | N | -0.45 |
| CTP1 | N | -0.44 |

| | | |
|---|---|---|
| YNL165W | N | -0.42 |
| MNL1 | N | -0.42 |
| EGT2 | N | -0.42 |
| KAP123 | N | -0.42 |
| HIS4 | N | -0.41 |
| MRH1 | N | -0.41 |
| POL1 | M | -0.4 |
| FAA3 | N | -0.4 |
| HSP150 | N | -0.4 |
| ASN1 | N | -0.4 |
| LYS12 | N | -0.39 |
| YHB1 | N | -0.38 |
| MCM2 | M | -0.37 |
| LEU9 | N | -0.37 |
| PSE1 | N | -0.36 |
| SEN34 | M | -0.35 |
| LYS21 | N | -0.35 |
| TPO4 | N | -0.34 |
| SSO1 | N | -0.34 |
| BAR1 | N | -0.33 |

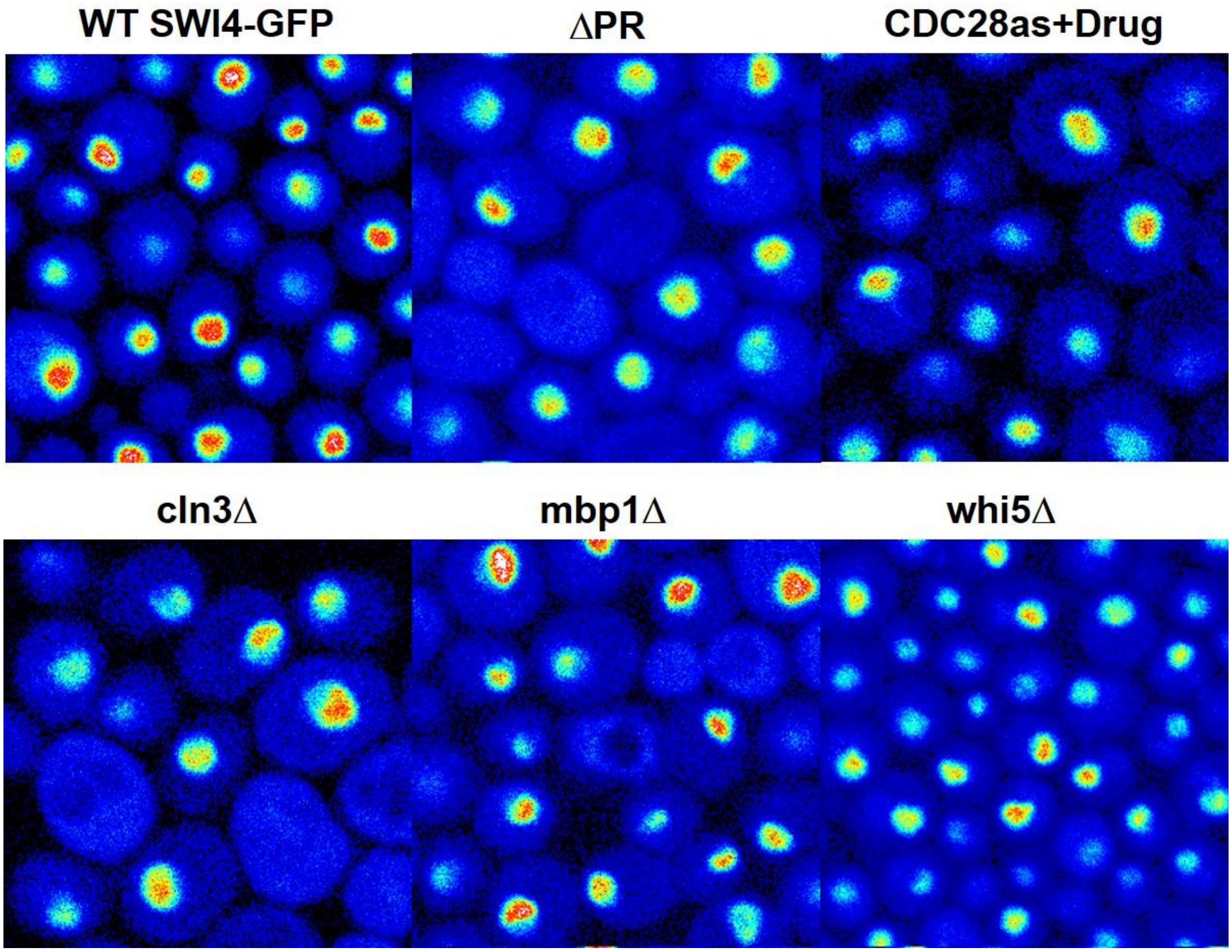

**Figure S1. Representative average images from sNB measurements.** Strains are indicated. Laser excitation intensity as identical in each experiment. Each average image is the result of 25 scans with a pixel dwell time of 40 μs. Image size scale is 20x20 μm. Intensity scales are similar. Nuclei were masked and the intensities within each nuclei were average over all nuclear pixels. Absolute number of GFP molecules in the excitation volume, N, was determined as the average intensity of the nucleus divided by the molecular brightness of GFP, $e_{GFP}$, calibrated each day using cells expressing GFP from a plasmid. The concentration was calculate by dividing N by the excitation volume, $V_{PSF}$, calibrated each day and Avogadro's number, $N_A$. Experiments were carried out as described by Dorsey et al 2018 (reference 6 in the main text).

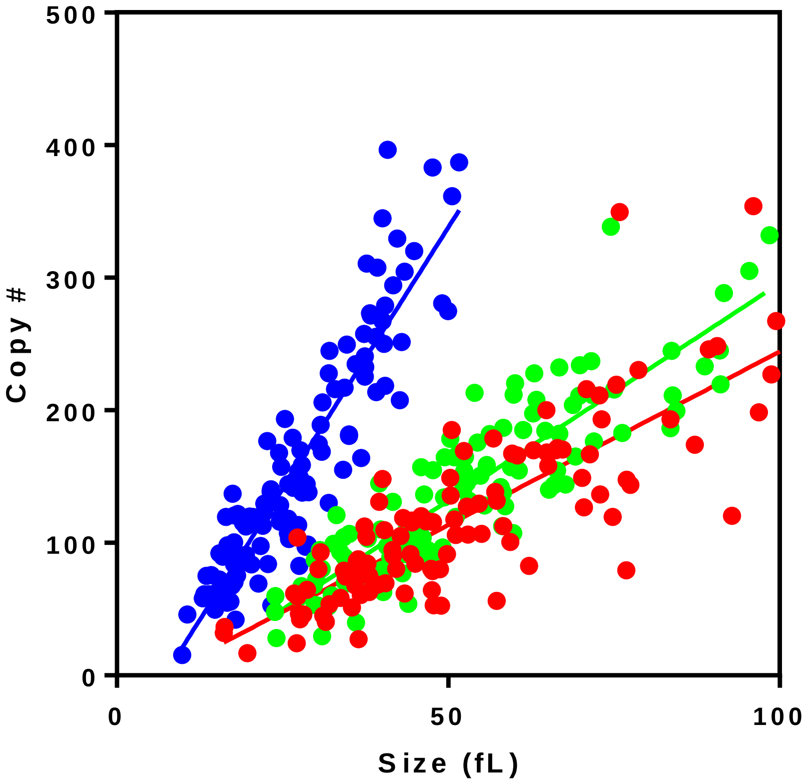

**Figure S2. Accumulation of Swi4-GFP vs size** . WT haploid (blue), WT diploid (green) and Diploid *ΔPR* (red) strains.

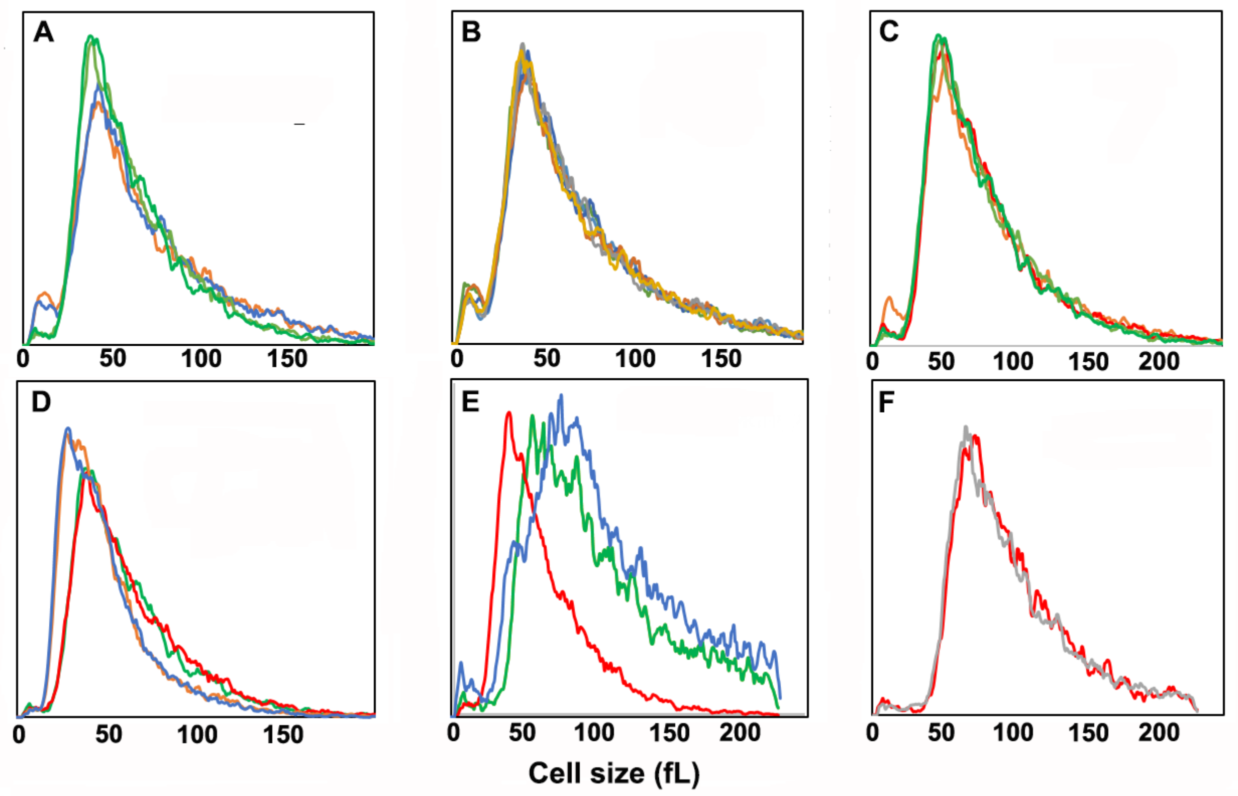

**Figure S3. Coulter counter cell size measurements.** A) Swi4-GFP (two replicates light and darker green) and Mut1 (two replicates, blue and orange), B) Mut1, Mut2 and Mut3 strains (two replicates each, Mut1+Swi4-GFP - dark blue and orange, Mut2+Swi4-GFP - gray and yellow, Mut3+Swi4-GFP - dark blue and orange, green and light blue), C) Swi4-GFP (two replicates – light and darker green) and *ΔPR*+Swi4-GFP (two replicates, red and orange), D) Swi4-GFP (two replicates – red and green) and *whi5Δ*+Swi4-GFP (two replicates – blue and orange), E) Swi4-GFP (red), cdc28as+Swi4-GFP – no drug (green) and *cln3Δ*+Swi4-GFP (blue) and F) Diploid Swi4:Swi4-GFP (gray) and diploid Swi4:*ΔPR*+Swi4-GFP (red).

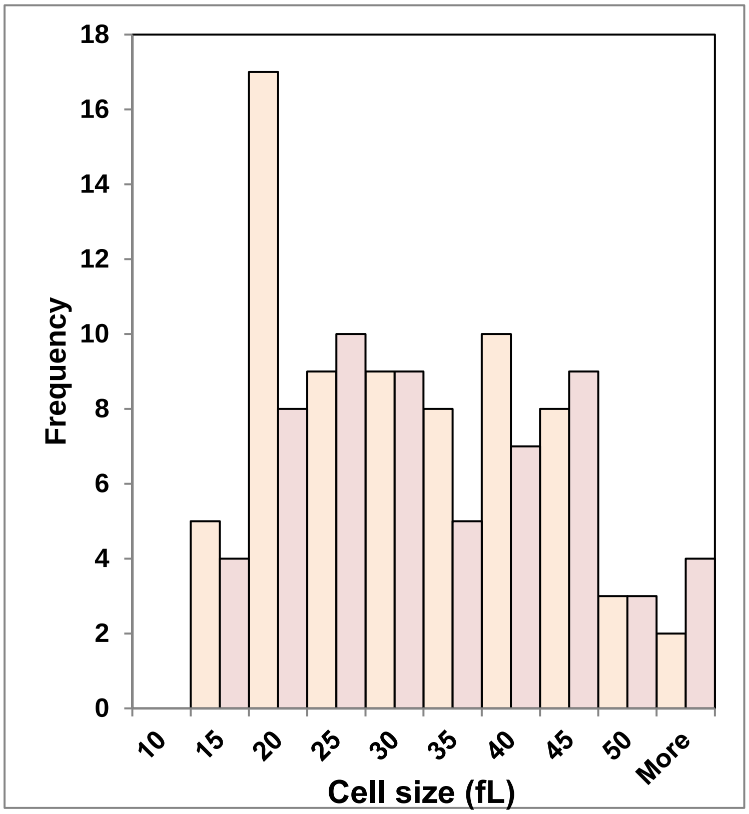

**Figure S4. Cell size histogram for the *ΔPR* strain.** Cell sizes were calculated for cells expressing nuclear Swi4-GFP. WT Swi4-GFP (peach), *ΔPR* Swi4-GFP (light pink).

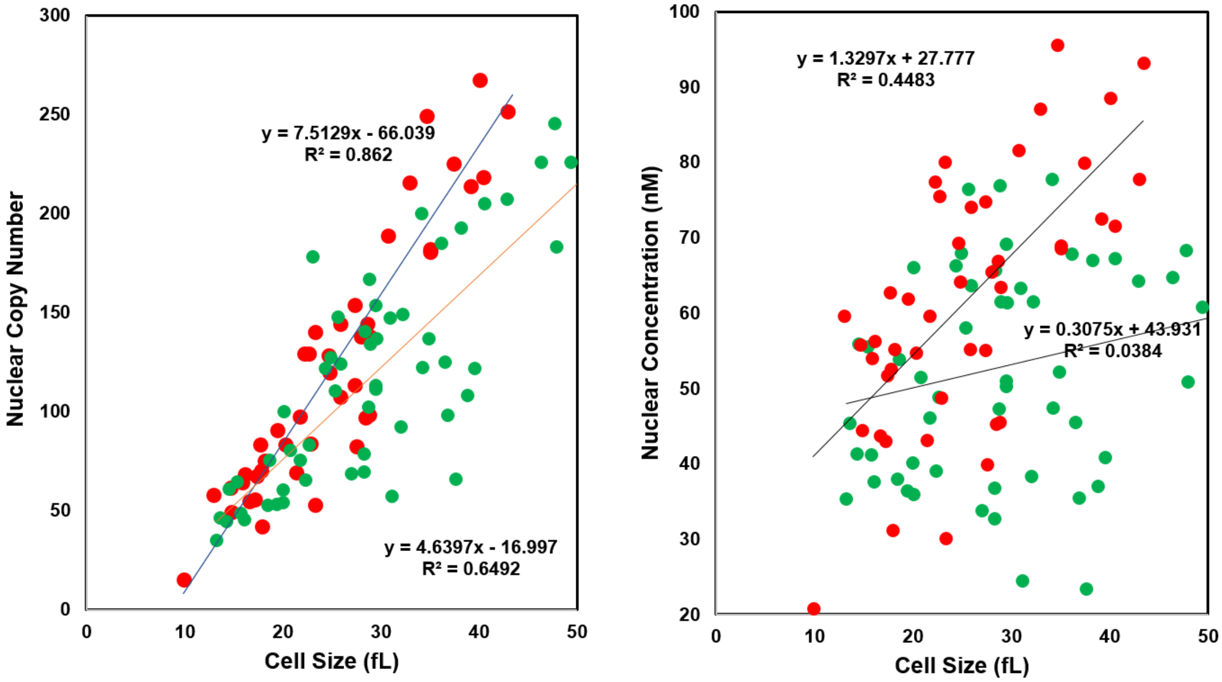

**Figure S5. The cell size dependence of Swi4 concentration exhibits larger cell-to-cell variation than the cell size dependence of Swi4 copy number**. A) Swi4-GFP nuclear copy number vs cell size for WT haploid cells expressing Swi4-GFP (red) and for *ΔPR* haploid cells expressing Swi4-GFP. B) A) Swi4-GFP nuclear concentration vs cell size for WT haploid cells expressing Swi4-GFP (red) and for *ΔPR* haploid cells expressing Swi4-GFP. Note the $R^2$ values are much higher for the copy number plots than for the concentration plots.

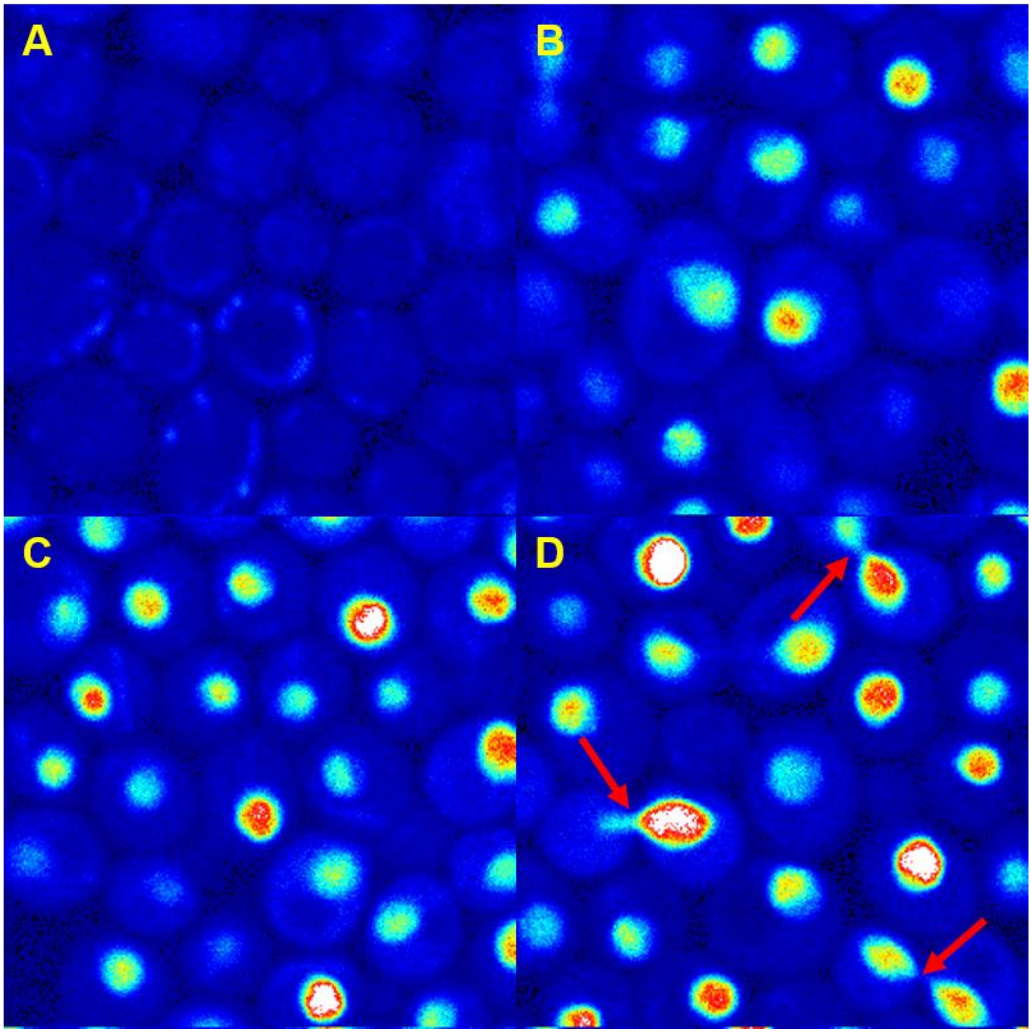

**Figure S6. BE2-dependent production of Swi4-GFP from the Z3EV promoter. A-D)** Images of FOV of the *Z3EV:Swi4-GFP/SWI4:Swi4-GFP* (MTy5189) strain in presence of 0, 2, 3, and 5 nM , respectively. Image full scale is 0-8 counts.
